# Biosynthesis of the azoxy compound azodyrecin from *Streptomyces mirabilis* P8-A2

**DOI:** 10.1101/2023.10.13.562208

**Authors:** Matiss Maleckis, Mario Wibowo, Tetiana Gren, Scott A. Jarmusch, Eva B. Sterndorff, Thomas Booth, Nathalie N. S. E. Henriksen, Christopher M. Whitford, Xinglin Jiang, Tue S. Jørgensen, Ling Ding, Tilmann Weber

**Affiliations:** The Novo Nordisk Foundation Center for Biosustainability, Technical University of Denmark, Kemitorvet, Building 220, 2800 Kgs. Lyngby, Denmark; Department of Biotechnology and Biomedicine, Technical University of Denmark, Kongens Lyngby, Denmark, Kemitorvet, Building 221, 2800 Kgs. Lyngby, Denmark; Singapore Institute of Food and Biotechnology Innovation (SIFBI), Agency for Science, Technology and Research (A*STAR), 138669, Singapore

## Abstract

Azoxy compounds are a distinctive group of bioactive secondary metabolites, characterized by a unique RN=N^+^(O^-^)R moiety. The azoxy moiety is present in various classes of metabolites that exhibit various biological activities. The enzymatic mechanisms underlying azoxy bond formation remain enigmatic. Azodyrecins are cytotoxic azoxy metabolites produced by *Streptomyces mirabilis* P8-A2. Here we cloned and confirmed the putative *azd* biosynthetic gene cluster through CATCH cloning followed by expression and production of azodyrecins in two heterologous hosts, *S. albidoflavus* J1074 and *S. coelicolor* M1146, respectively. We explored the function of 14 enzymes in azodyrecin biosynthesis through gene knock-out using CRISPR-Cas9 base editing in the native producer, *S. mirabilis* P8-A2. The key intermediates were analyzed in the mutants through MS/MS fragmentation studies, revealing azoxy bond formation via the conversion of hydrazine to azo compound; followed by further oxygenation. Additionally, *N*-oxygenase and dehydrogenase activities were confirmed among 8 core biosynthetic genes and five helper genes. Moreover, the distribution of the azoxy biosynthetic gene clusters across *Streptomyces* spp. genomes is explored, highlighting the presence of these clusters in over 20% of the *Streptomyces* spp. genomes and revealing that azoxymycin and valanimycin are scarce, while azodyrecin and KA57A like clusters are widely distributed across the phylogenetic tree.

## Introduction

Azoxy compounds are a group of intriguing bioactive molecules sharing the azoxy moiety (RN=N^+^(O^-^)R)^1^. Their diverse biological activities and unique chemical structures position them as an important class of metabolites. *Streptomyces* are known to be prolific producers of azoxy compounds, such as elaiomycins^2–9^, LL-BH872α^10^, valanimycin^11,12^, KA57A^13^, maniwamycins^14–16^, jietacins^17^, azodyrecins^18,19^, azoxymycins^20^, O-alkylazoxymycins^21^, DC-8118 A-B^22^, geralcin C^23^ and an unnamed azoxy compound^24^. The latter azoxy compound was identified through heterologous expression of azoxy biosynthetic gene cluster (BGC) from *S. avermitilis* MA-4680 in *S. coelicolor* M1152 and *S. lividans* TK24. The new molecules were detected in the heterologous host and exact mass suggested them to be of azoxy origin, while the structure of the unnamed compound is yet to be elucidated. In view of the various biological activities of azoxyl compounds, such as antibacterial valanimycin and DC-8118, antifungal maniwamycins, KA57A and O-Alkylazoxymycins and cytotoxic azodyrecins, jietacins, elaiomycins and geralcin C, understanding the mechanism behind the biosynthesis of these compounds will unlock novel applications in medical, agriculture, dye, and other industries^25^.

Comparative analysis of the structure of natural azoxy compounds has led to proposition of two distinct routes for the azoxy biosynthesis^25^. One route involves dimerization for assembly of azoxymycins and O-alkylazoxymycin. This was confirmed through pioneering work into the azoxymycin‘s azoxy functional group formation through a radical-based coupling^26^ and suggested a combination of enzymatic and non-enzymatic steps^26^ in the azoxy bond assembly. The other proposed route links two different subunits such as two different amino acids, fatty acid and amino acid, or polyketide synthase derived subunit and amino acid and is the case for remaining of the known *Streptomyces* spp. azoxy compounds, Figure 1.

**Figure 1:**
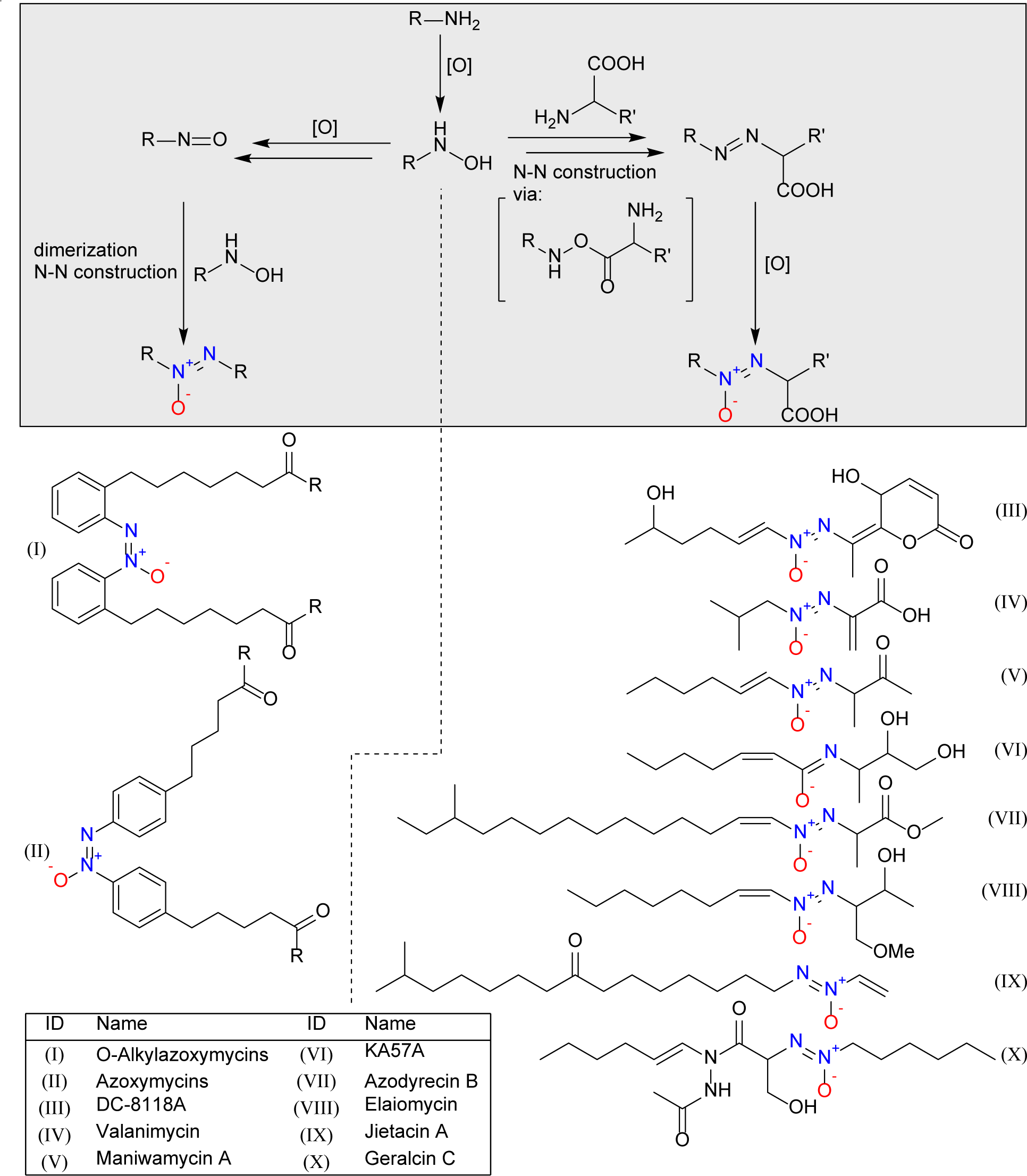
Proposed two routes for azoxy compound biosynthesis adapted and modified from Wibowo, M and Ding, L, 2020^25^, where one assembly path utilizes dimerization, while the other one cross links with carboxylic group, followed by rearrangement. The assigned route for the biosynthesis of *Streptomyces* spp. azoxy compounds is proposed, where only O-alkylazoxymycins (I) and azoxymycins (II) belong to the first route.

Previous *in vitro* enzymatic studies and feeding experiments^27–37^ related to valanimycin biosynthesis have provided insights into the almost complete biosynthesis pathway, with established function of eleven out of fourteen genes responsible for the assembly of valanimycin^29,34^. Valanimycin is assembled from two amino acids, valine and serine. The valine gets decarboxylated^34^ by VlmD and subsequently hydroxylated^34^, by VlmH and VlmR, and attached to serine^36^ by VlmA, whereafter the molecule undergoes re-arrangement resulting in the formation of an azoxy bound through an unelucidated process. At the very end, the valanimycin hydrate is phosphorylated^37^ followed by dehydration^37^ by VlmJ and VlmK. The three genes^34^, *vlmB*, *vlmG* and *vlmO* remain enigmatic and are hypothesized to play a role in azoxy bond formation.

Azoxymycins A-C are produced by *S. chattanoogensis* L10^20^ and their biosynthesis was confirmed by gene deletion and verified by three mutants^20^ *ΔazoFG*, *ΔazoJ* and *ΔazoC*. Through in vitro characterization of AzoC^26^, the azoxy bond formation in azoxymycins is a combination of enzymatic and non-enzymatic coupling cascade reaction. AzoC encodes nonheme diiron N-oxygenase that oxidizes the amine to a nitroso group, which allows for dimerization into azoxymycins through non-enzymatic reactions facilitated by redox coenzyme pairs^26^.

Recently isolated from *S. mirabilis* P8-A2, azodyrecins A-C^18^ represent a set of aliphatic azoxy metabolites. In this study, we selected azodyrecins as the model to uncover the biosynthesis of aliphatic azoxy compounds. Here, we confirm the biosynthetic gene cluster (BGC) responsible for azodyrecin biosynthesis through cloning and heterologous expression in *S. coelicolor* M1146^38^ and *S. albidoflavus* J1074^39^. Furthermore, by constructing knock-out (KO) strains targeting fourteen genes within the BGC, we gained insights into essential genes for the azodyrecin biosynthesis. The KO in *S. mirabilis* P8-A2 was achieved using CRISPR-Cas9 base editing tool, CRISPR-BEST^40,41^, which was used to introduce a stop codon in the upstream region of target gene coding sequence without creation of double stranded break and risk of genome rearrangement. Through metabolomic guided analysis of resulting strains we were able to propose the function of certain genes within azodyrecin biosynthesis pathway. Lastly, our exploration of the prevalence of various azoxy biosynthetic gene clusters across the phylogenetic tree of medium- and high-quality *Streptomyces* spp. genomes offers valuable insights into the compounds’ potential significance within the lifestyle of *Streptomyces*.

## Results and Discussion

### Genome assembly of azodyrecin producer *Streptomyces* sp. P8-A2

Whole genome sequencing of the azodyrecin-producing strain was performed using Oxford Nanopore and Illumina sequencing. The assembly revealed a chromosome of 11.468.629 bp with a GC content of 70%. Five additional scaffolds were assembled, of which three contain *oriC* region, predicted using DoriC 12.0 database^42^ search, indicating that they are separate entities of linear/circular plasmids. The assambled sequence was predicted to contain 40 biosynthetic gene clusters using antiSMASH 7.0.1^43^, relaxed detection strictness. The strain was identified as *Streptomyces mirabilis* by GTDB-Tk (v2.1.1) with an ANI% of 96.64 to GCF_014650275.1^44^.

### Azodyrecin biosynthetic gene cluster of *Streptomyces mirabilis* P8-A2

Production of azodyrecins have been reported in two other *Streptomyces* sp. by recent work of Choirunnisa, A. R. et al.^19^, where they elucidated tailoring step of methylation by S-adenosyl methionine (SAM) dependent methyltransferase, Ady1, in vitro and proposed BGC for azodyrecin biosynthesis in two azodyrecin producer strains. One of the reported BGCs in *Streptomyces* sp. A1C6 (NCBI GenBank: LC712331) is highly similar to *S. mirabilis* P8-A2, sharing average nucleotide identity (ANI) of 95.7% and gene protein identities of 81 to 98 %. Interestingly, the BGC from *Streptomyces* sp. RM72 (NCBI GenBank: LC712332) is significantly distant from *S. mirabilis* P8-A2, ANI of 60 % and protein sequence similarity for 10/12 protein sequences between 32 and 75 %, while still producing azodyrecins A-F.

The proposed azodyrecin BGCs have not been experimentally validated and therefore we began our analysis by assessing the BGC boundaries. First, we compared the BGCs of all known producers of azoxy compounds in *Streptomyces* spp. (Table S.1) and thereafter analyzed antiSMASH^43^ annotated gene function predictions to determine the BGC borders. BGCs encoding the biosynthesis of azoxy-compounds have been confirmed for azoxymycin^26^, valanimycin^34^ and unnamed azoxy compounds of *S. avermitilis* MA-4680^24^. Furthermore, BGCs have been proposed for KA57-A^45^ and azodyrecins without validation by KOs or heterologous expression. For azoxymycins, only biosynthetic gene sequences have been published. Searching for all of the described genes, we identified a closely related 14kb BGC region in the *S. chattanoogensis* NRRL ISP-5002 sharing 97.9 % - 99.6 % nucleotide and 96.2 % - 100.0 % amino acid identity (Table S.2). No genetic information is available for the remaining seven known azoxy compounds produced by *Streptomyces* spp. Using the genomic information, we generated an alignment of BGCs and identified the genes shared between them using clinker^46^ (Figure 2, characterized valanimycin gene functions:Table S.3). As already indicated by the overall ANI analysis, the azodyrecin BGCs of *S. mirabilis* P8-A2 and *S.* sp. A1C6 are highly similar and syntenic, whereas the BGC of *Streptomyces* sp. RM72 shows lower similarities and a different gene organization. BGCs encoding other azoxy-compounds, i.e., valanimycin, KA57-A and azoxymycin share eight genes within the proposed azodyrecin BGC, five of which are shared across all azoxy BGCs, except for azoxymycin. As the upstreamand downstream regions of the proposed core biosynthesis genes encoded further transporters, regulators, cytochrome P450 and other activities, we included seven additional genes upstream and thirteen downstream compared to already published BGCs, Figure 2 highlighted BGC. The sequence of the azodyrecin BGC from *S. mirabilis* P8-A2 is deposited in MiBIG database under accession number BGC0002805 and detailed overview is presented in Figure 4.

**Figure 2:**
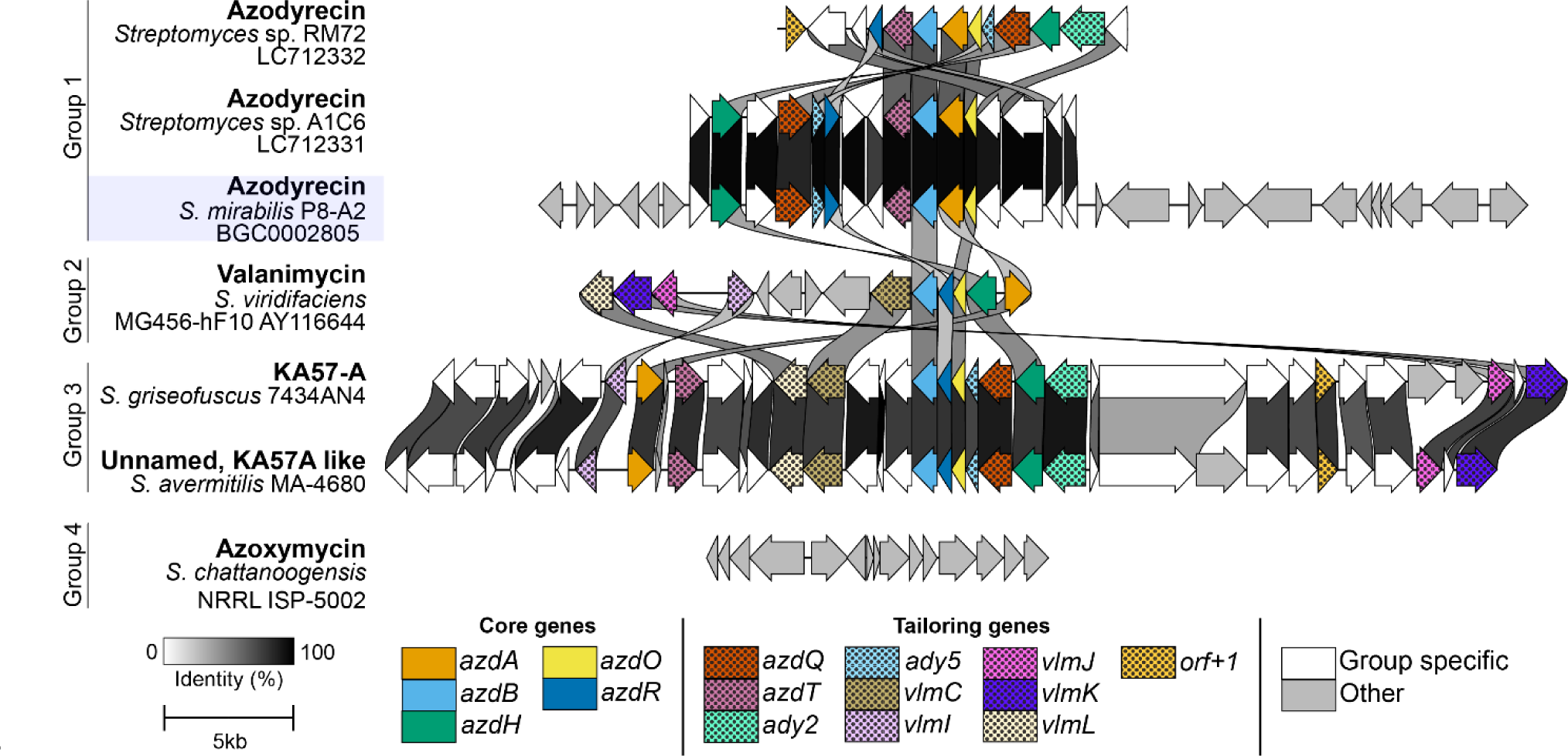
Gene cluster comparison using clinker^46^ of known and proposed biosynthetic gene clusters (BGCs) for azoxy compounds of *Streptomyces* spp. The BGCs are grouped based on their cluster similarity. The colors reflect if the gene is group specific or shared with other groups. Tailoring genes are shared across two groups, while core genes are shared across all, except group 4. The *S. mirabilis* P8-A2 azodyrecin BGC is highlighted and visualized at proposed size.

### Production of azodyrecin in heterologous hosts

To confirm the identity of the *azd* BGC, it was cloned and heterologously expressed in *S. albidoflavus* J1074^39^ and *S. coelicolor* M1146^38^, using a modified version^47^ of the CATCH-cloning^45^ procedure. This method involves the targeted extraction of the predicted *azd* BGC using CRISPR-Cas9 and single guide RNAs (sgRNAs) in an in vitro system. The extracted *azd* BGC was subsequently inserted into the shuttle vector pXJ157 and then transferred into the heterologous host via biparental conjugation. Production of azodyrecin was confirmed by LC-MS analysis by comparison of azodyrecin B standard to culture extracts of the heterologous expression hosts (Figure 3). Production levels were similar with slightly higher production detected by *S. coelicolor* M1146. The products were verified by MS/MS spectra comparison, which confirmed their identity (Figure S.8). The successful detection of azodyrecin in heterologous hosts confirms that the candidate cluster contains all of the required genes for the azodyrecin biosynthesis.

**Figure 3:**
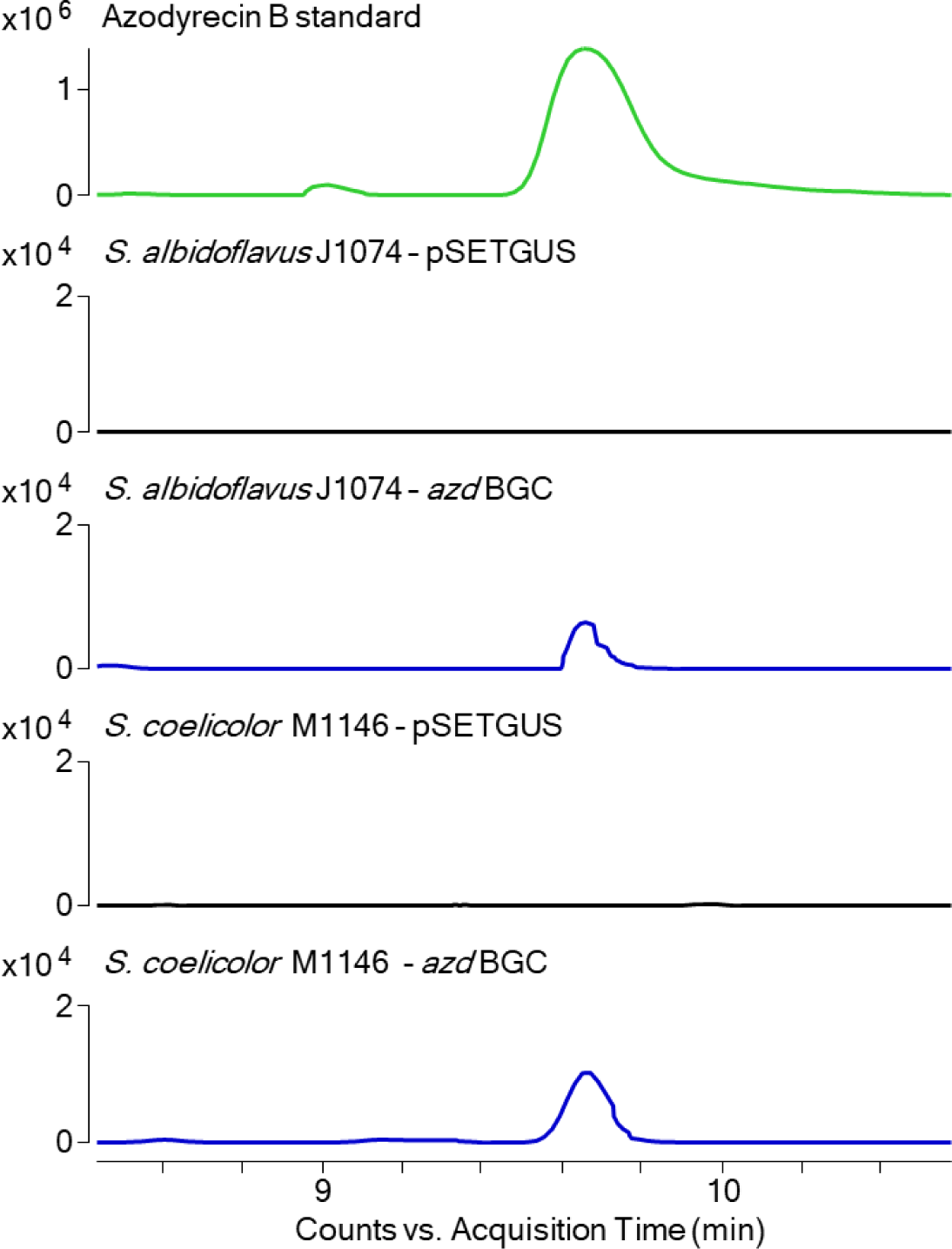
Extracted ion chromatogram (EIC) of azodyrecin B (m/z 341.2804 [M+H]^+^), in standard and extracts from the heterologous expression host with *azd* BGC integrated compared to negative control, pSETGUS^48^.

**Figure 4:**
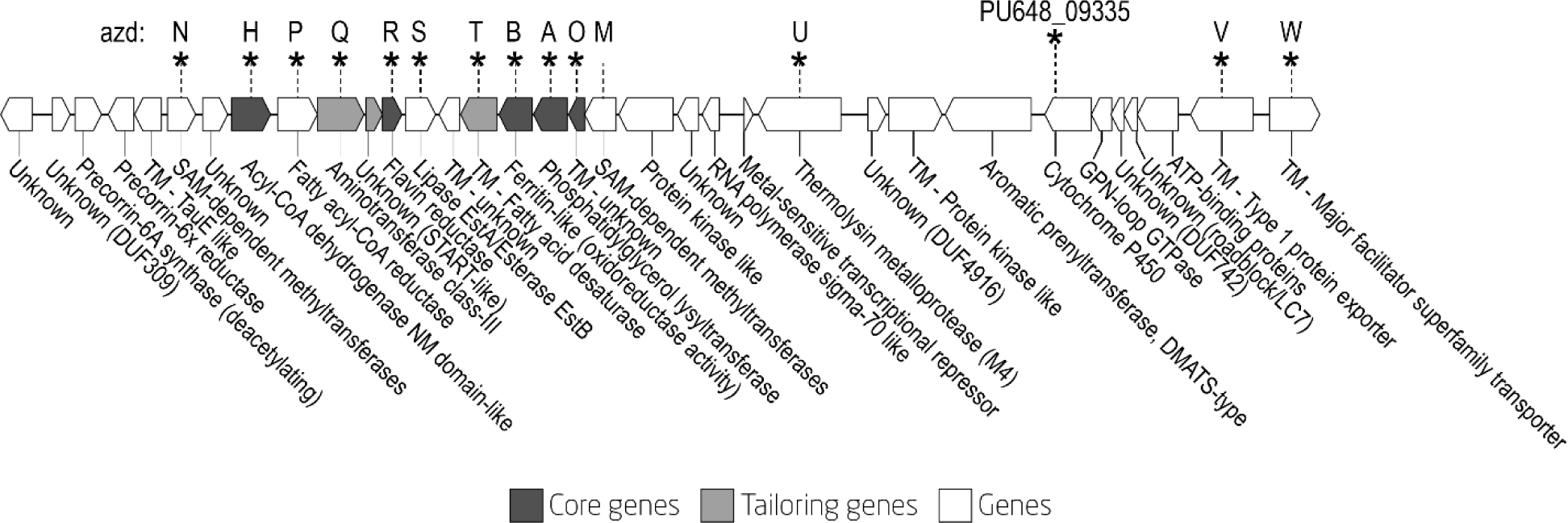
Overview of *azd* biosynthetic gene cluster. Genes that were studied in this work are labelled with a star and their assigned gene names. The putative functions were predicted with InterPro^49^ scan v. 5.63-95.0. TM: transmembrane.

### Insights into the azodyrecin biosynthesis through knockout strain analysis

The pathway for biosynthesis of azodyrecins is largely unknown. Only the methylation of the carboxylic group by *azdM* has been experimentally confirmed^19^. To investigate the role of the individual key biosynthetic genes in the *azd* BGC, we established CRISPRBEST^40^ base editing in *S. mirabilis* P8-A2. We successfully generated fourteen KO strains by introducing a stop codon in the N-terminal region of the CDS, which prevents the synthesis of mature protein (Figure 4; Table S.5). The targets for KO study were selected based on their conservation across azoxy BGCs (Figure 4 core and tailoring genes) and genes encoding predicted function that might be involved in upregulation azodyrecins precursor availability, dehydration of fatty acid, transport, or other aspects of biosynthesis, but are not shared across other known azoxy BGCs.

We applied a feature based molecular network (FBMN)^50^ to analyze the untargeted metabolomic data and map precursors and intermediates in the different KO strains. Through network analysis, we compared the presence and abundance of ionized masses of WT culture extracts to the mutant. This way the data allowed us to propose 11 intermediates and to which we proposed their structures based on exact mass and fragmentation analysis (Figure 5. A, Figure S.9 – Figure S.15). Comparison of LC-MS data showed that compounds **13**-**17** were not detected in the WT strain, but could be seen in *azdT*^STOP^*, azdB*^STOP^*, azdH*^STOP^*, azdR*^STOP^ and *azdS*^STOP^ mutants, respectively (Figure 5.B, Figure S.16 – Figure S.21). Production of azodyrecins A-C (**1-3**) was abolished in *azdT*^STOP^*, azdB*^STOP^ and *azdH*^STOP^ mutants, although they still produced several intermediates, **4-15**, **10-15** and **16-17**, respectively. Azodyrecins were still produced in *azdR*^STOP^*, azdS*^STOP^*, azdU*^STOP^*, azdN*^STOP^ and *azdW*^STOP^ mutants, however, intermediate compounds were detected at different relative abundances compared to the WT, suggesting that they are involved as helper genes but are not required for the biosynthesis of azodyrecins. The *PU648_09335*^STOP^ mutant, in which a putative cytochrome P450 was inactivated, still produced azodyrecin and thus seems not to be involved in azodyrecin biosynthesis. In all other mutants we did not detect any of the compounds (**1-17**) in the LCMS analyses.

**Figure 5:**
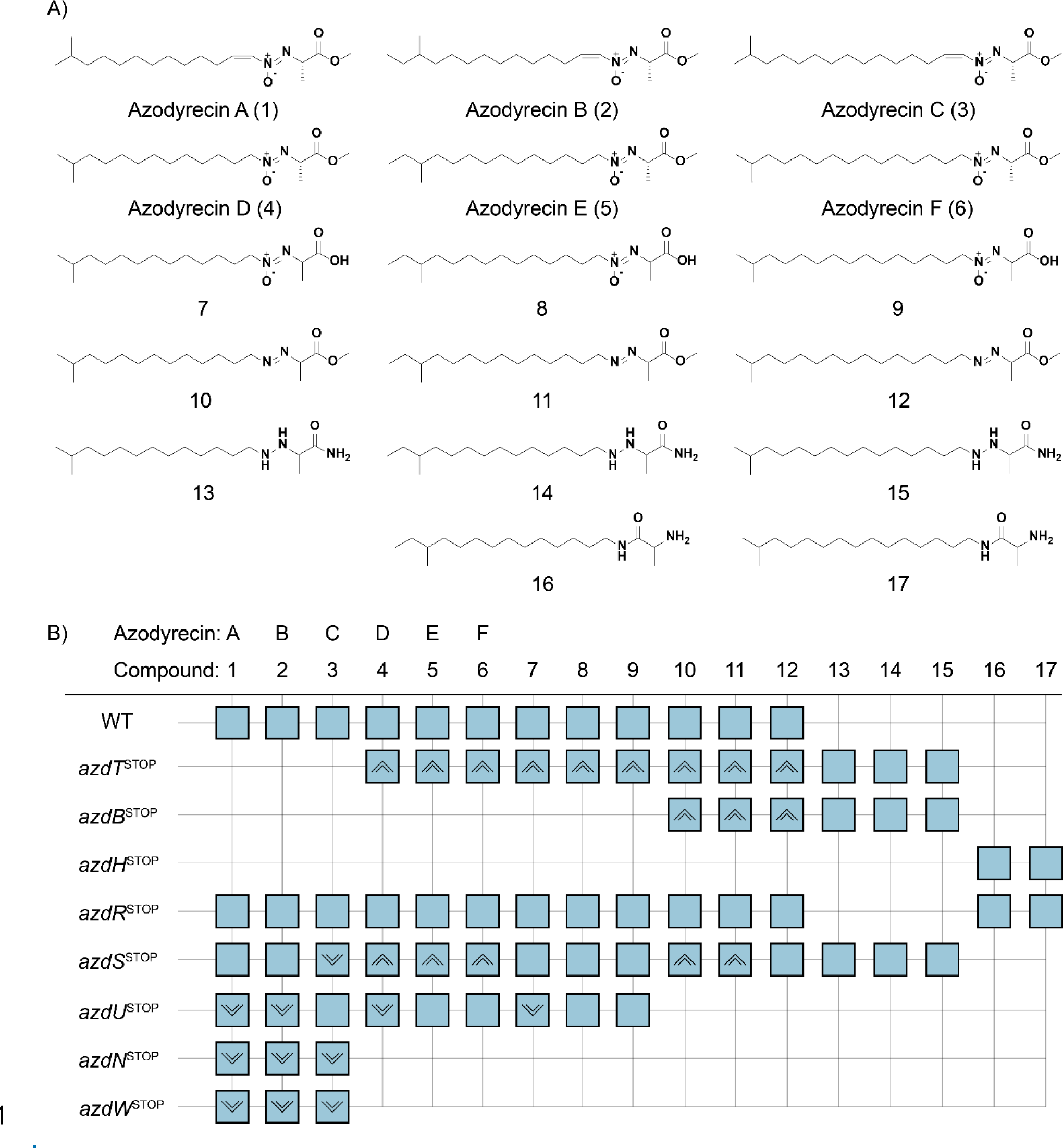
**A**) Overview of compounds **1-17** detected in the metabolomics data of the WT and/or knock out strains. Azodyrecins A-F (**1**-**6**) are known structures, while **7**-**17** are proposed structures based on exact mass, MS/MS studies, and comparison with the known compounds **1**-**6** that were found in the azodyrecin knockout strains. **B**) Presence/absence matrix of LC-MS detected compounds in specific strains is indicated with a square; arrows indicate relative amount compared to the wild type strain, four times higher or lower.

*azdH* and *azdR* exhibited similarity to *vlmH* ^28,34^ and *vlmR*^27,34^, which are responsible for the *N*-hydroxylation of isobutylamine^34^ in valanimycin biosynthesis (Table S.2). Inactivation of either of these genes results in the formation of compounds **16-17**, with an amide bond formed through linkage of fatty-amine to the activated alanine. In the *azdH*^STOP^ mutant we did not detect downstream intermediates, only compound **16-17**. The *azdR*^STOP^ mutant produced same compounds as the WT strain and additionally compounds **16-17** (Figure S.21). Based on these findings we suggest that *azdH* is responsible for *N*-hydroxylation of the fatty amine, while *azdR* inhibits formation of this off-product and directs flux towards *N*-hydroxylation. It is likely that the formation of compounds **16-17** is due to unspecificity of AzdA in absence of AzdR, allowing to deliver alanine to amine. *azdA* encodes a putative phosphatidylglycerol lysyltransferase, a family of enzymes, which are known to facilitate transfer of an amino acid from AA-tRNA to a membrane bound substrate. It has been discovered in pathogens such as *Staphylococcus aureus* where the enzyme modifies phosphatidylglycerol into lysylphosphatidylglycerol to modify membrane charge and avoid detection by defensins. In the biosynthesis of azodyrecin, we propose that AzdA, in the presence of AzdR, specifically transfers alanine from tRNA^Ala^ to a hydroxylated fatty amine. Conversely, in the absence of AzdR, AzdA is also capable of transferring alanine from tRNA^Ala^ to a non-hydroxylated fatty amine . The *azdO* gene encodes a 155 AA transmembrane protein with three predicted transmembrane helices according to InterPro^49^ scan v. 5.63-95.0. The gene does not have any characterized homologues and therefore function cannot be predicted. No intermediates could be detected in the *azd*A^STOP^ and *azd*O^STOP^ mutants, which could be due to the instability of the intermediates. AzdO might facilitate recruitment of the substrate and other catalytic enzymes, or even be responsible for rearrangement and formation of hydrazine and azo bond in the proposed intermediates, however it is unclear which reactions are spontaneous and which are catalyzed by enzymes at this stage.

The key finding was the function of AzdB, that we identified to be responsible for azoxy bound formation. The *azd*B^STOP^ mutant produced hydrazine **13-15** and azo compounds **10-12**. We did not observe oxidation of these hydrazine and azo compounds in the mutant, suggesting that the AzdB is required to produce the azoxy group and carries out oxidation reaction of hydrazine/azo group. This evidence confirmed that *azdA*, *azdB*, *azdH* and *azdO* code for the enzymes carrying out the essential functions for the azoxy bond formation. These genes are conserved in a group of azoxy compounds where azoxy bound is formed through N-N constitution followed by nitrogen oxidation, i.e., azodyrecins, KA57A, maniwamycins, valanimycin, eliomycins and others.

We also targeted other genes that might be involved in the biosynthesis of azodyrecins, their transport and other aspects of biosynthesis. The production of azodyrecins A-C was inhibited in the *azdT*^STOP^ mutant, while intermediates **4-12** were produced at higher abundances compared to the WT. This indicates that AzdT is a fatty acid desaturase and is responsible for double bond formation in the fatty chain moiety of azodyrecins. Desaturation of the fatty amine was only observed in the final products of the pathway, post methylation, suggesting that AzdT performs the final reaction converting azodyrecin D-F (**4-6**) into azodyrecin A-C (**1-3**). All of the intermediates were produced in higher abundance in the *azdT*^STOP^ compared to the WT and a similar increase was also observed in the *azdS*^STOP^ mutant. However, the *azdS*^STOP^ mutant was still able to produce azodyrecin A-C (**1-3**). According to the ESTHER database^51^*, azdS* encodes a putative alpha/beta fold hydrolase with predicted activity belonging to lipase class 2^50^ and it is unclear what role this gene has in resulting in relatively high levels of intermediates (**4-6**).

The fatty acid moiety in azodyrecins is an iso-fatty acid, synthesized using a variety of different amino acid as starter units. Starter units must be deaminated and prolonged by malonate chain extension, by a fatty acid synthase not encoded in the BGC.^52^ Therefore the azodyrecin B (**2**) starter unit is isoleucine, while for azodyrecin A and C (**1, 3**) the starter unit should be valine with difference of one malonate chain extension. These iso-fatty acids could originate from primary metabolism and used in biosynthesis, as they help to control membrane fluidity in bacteria^52^ and their abundances are species specific^53–55^. The *azdQ* gene encodes a putative aminotransferase, which could both aminate the fatty acid. It is unclear whether the fatty acid is aminated following decarboxylation or reduction of the carboxy group into an alcohol. It is likely that the latter hypothesis is true, as *azdP* encodes a putative alcohol forming fatty acid reductase, which could be responsible for the formation of an iso-fatty alcohol, which thereafter is aminated by *azdQ*. Such iso-fatty amines have previously been described in *Streptomyces* sp. NK14819 strain and were named medelamines A and B^56^ and could serve as the precursors in the biosynthesis of azodyrecins A-B (**1-2**). The ABC-transporter, AzdV, appeared to be essential for biosynthesis, as none of the compounds **1-17** were detected in the KO mutant. Its role might be transport of biosynthetic enzymes to the periplasmic space where azodyrecins might be synthesized, however it could also be toxicity requiring the cell to stop its production of azodyrecins. A mutant in *azdW* encoding a major facilitator superfamily transporter KO mutant produced only azodyrecins A-C and at low levels. The M4 metalloproteases, *azdU*^STOP^, and methyltransferase, *azdN*^STOP^, mutants also produced azodyrecins A-C at reduced amounts, but it is not clear what exact function these genes have in the pathway. Using the acquired data from the KO mutants we can propose an azodyrecin biosynthesis pathway (Figure 6).

**Figure 6:**
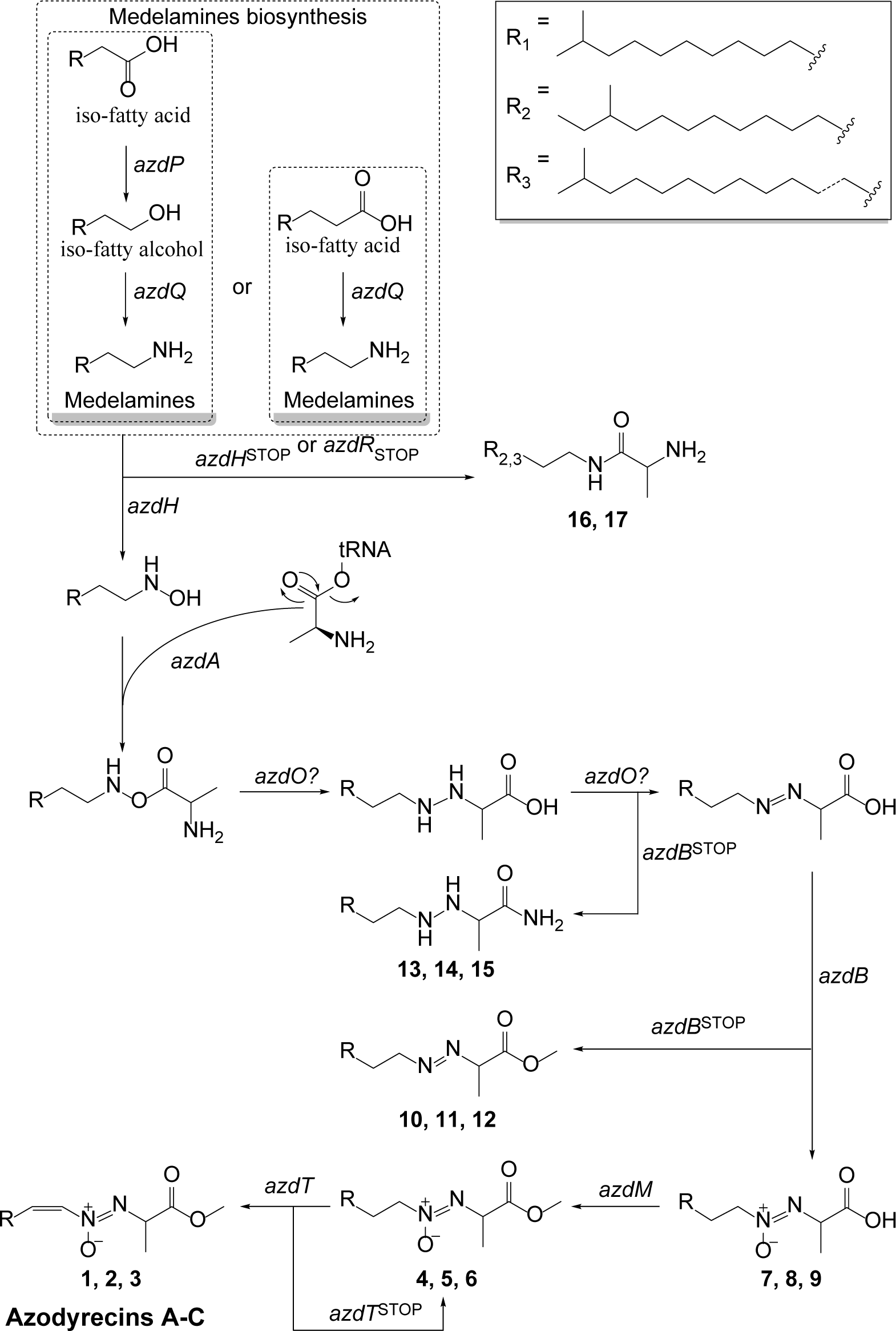
Hypothetical pathway for azodyrecin biosynthesis where the intermediates detected in the KO mutants are indicated with numbers.

### Distribution and abundance of azoxy biosynthetic gene clusters across phylogeny **of *Streptomyces* sp.**

With new knowledge about azoxy compound biosynthesis, we investigated the distribution of such pathways in the *Streptomyces* genus. We mapped known BGCs of azoxy compounds to a genome scale phylogenetic tree of 1528 *Streptomyces* medium/high quality genomic sequences (Figure 7) acquired from NCBI. We identified that large number of sequenced *Streptomyces* spp. to contain azoxy BGC, 351 out of the 1535 (22 %) genomes, while only 16 (1 %) of the genomes contained more than one azoxy BGC.

**Figure 7:**
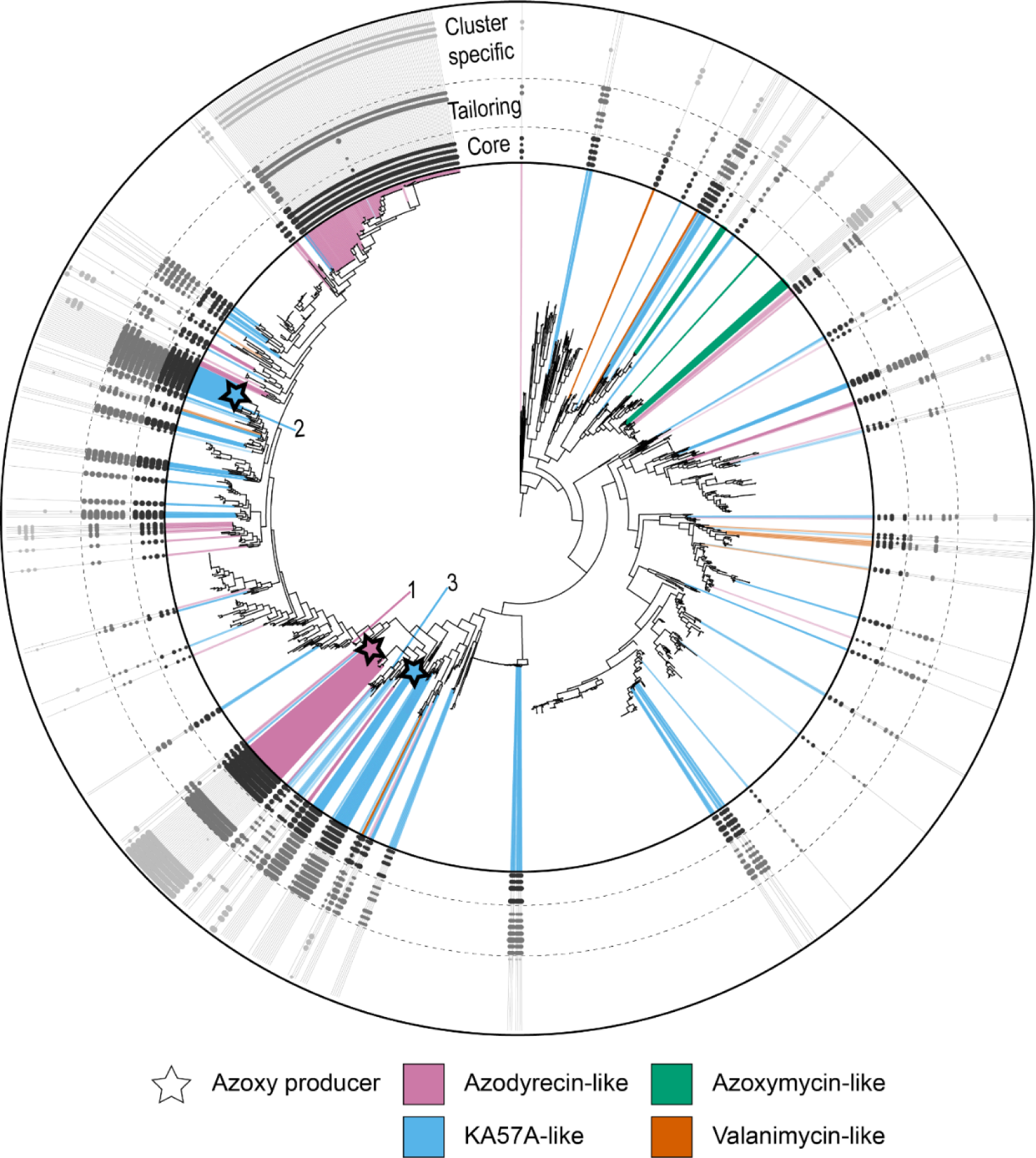
*Streptomyces* spp. azoxy BGC distribution across WGS phylogenetic tree *of Streptomyces spp.* generated using autoMLST^57^, including presence absence matrix of key biosynthetic genes for these compounds. The inner circle contains phylogenetic tree highlighting genomes containing BGCs similar to one of the four input clusters and are colored based on similarity to the known azoxy BGCs, while the intensity of the color reflects on the similarity the linked BGC. The WGS nodes of known producers are highlighted with a star: (1) *S. mirabilis*, (2) *S. griseofuscus* and (3) *S. avermitilis.* The outer circle shows presence/absence matrix of defined core, tailoring and cluster specific proteins. The size of the dots in the outer circle represents gene AA-identity to the input cluster protein sequence.

To have a closer look into distribution of different azoxy BGC types, we created four groups, azodyrecin-like, valanimycin-like, KA57A-like and azoxymycin-like. The grouping reflected on the BGC similarity and their chemical structure moiety differences. Azodyrecin-like compounds are composed of iso-fatty and amino acid moieties, valanimycin-like from two amino acids, KA57A-like from short chain fatty and amino acid, and the azoxymycin-like compounds by dimerization of phenolic monomers. The grouping is visualized in Figure 2 where shared genes between the different groups are highlighted.

The results showed that the valanimycin-like and the azoxymycin-like BGCs were identified in 14 and 12 genomes respectively. The azoxymycin-like BGCs appear to be clade specific while valanimycin-like BGCs are identified in distant clades across the tree. BGCs encoding azodyrecin-like and KA57A-like compounds are most abundant, detected in 191 (12 %) and 136 (9 %) genomes respectively. While azodyrecin-like BGCs were the most abundant, many originate from closely related strains indicating a sampling bias in our dataset. Interestingly, we detected an azodyrecin-like BGC in the model streptomycetes *S. coelicolor* A3, although no azoxy compounds were reported in this strain. We compared this and other randomly selected azodyrecin-like BGCs using clinker^46^ and could verify that the genes are clustered in all the BGCs (Figure S.22). These BGCs likely encode new undiscovered azoxy compounds that bear similarity to azodyrecins.

The detection of KA57A-like BGCs showed slightly lower numbers, however it appears that this group is most spread across different clades. The distribution of KA57A-like BGCs could relate to higher diversity in this group of compounds. Previously structure similar azoxy compounds such as elaiomycins and maniwamycins have been discovered and there is potentially larger chemical structure variance for this group compared to azodyrecin like compounds. In recent work by Tanaka et al. 2023, they discovered analogues of KA57A in the same producer strain. These compounds were biosynthesized using different amino acids, valine and isoleucine instead of serine, however, they were azides and not azoxy compounds. Considering that the BGC is scattered across many clades and the relative size of the BGC, it is likely that similar compounds to KA57-A will be discovered in S*treptomyces* sp. For the strains in which we found azoxy BGC, we plotted the genes that were identified to be contained within the cluster, to have a better understanding of what is contained within the cluster and if the cluster is a hybrid of two.

The rules used to identify the four types of azoxy BGCs have been wrapped into antiSMASH rules allowing everyone to detect azoxy clusters in the future release.

## Methods

### Bacterial cultures and cultivation

Microorganisms used in the study were *Escherichia coli* ET12567/pUZ8002^58,59^, *Escherichia coli* ET12567/pUB307^58,60^, *Escherichia coli* Mach1 (Thermo Fisher Scientific; C862003) and *Escherichia coli* BAC-Optimized Replicator v2.0 (Lucigen; 60210–1), *Streptomyces mirabilis* P8-A2^18^, *Streptomyces coelicolor* M1146^38^ and *Streptomyces albidoflavus* J1074^39^.

Cultivation of *E. coli* strains was performed at 37 °C in LB (20g/L LB Broth (Lennox) (Sigma-Aldrich; L3022), 2xYT (tryptone 16g/L (Millipore; T9410), yeast extract 10g/L (Thermo Fisher Scientific; LP0021B), sodium chloride 5g/L(VWR; 470302)) or S.O.C. (tryptone 20g/L (Millipore; T9410), yeast extract 5g/L (Thermo Fisher Scientific; LP0021B), sodium chloride 0.5g/L(VWR; 470302)) medium. *Streptomyces* spp. were cultured at 30 °C in SFM^61^ (soya flour 20g/L (fettreduziert Bio Sojamehl; Hensel, Germany)), D-mannitol (Sigma-Aldrich; M4125)) or ISP2 (yeast extract 4g/L (Thermo Fisher Scientific; 212750), malt extract 10g/L (Sigma-Aldrich; 70167), glucose 4g/L (Sigma-Aldrich; G7021)). For agar plates the media was prepared with 2% w/v of agar (Sigma-Aldrich; 05040). For conjugations, SFM media was supplemented to contain final concentration of 10 mM MgCl2 (Sigma-Aldrich; M1028). When selection was required, the medium was supplemented with following antibiotics and their final concentrations: 100 µg/mL apramycin sulfate (Sigma-Aldrich; A2024), 25 µg/mL chloramphenicol (Sigma-Aldrich; C0378), 50 µg/mL kanamycin sulphate (Sigma-Aldrich; K1377) and/or 25 µg/mL nalidixic acid (Sigma-Aldrich; N8878).

### DNA isolation and sequencing of *Streptomyces mirabilis* P8-A2 genome

Sequencing of *Streptomyces* sp. P8-A2 was performed using both Oxford Nanopore and Illumina. The DNA extraction for Oxford Nanopore sequencing was done according to protocol by Alvarez-Arevalo et al. 2023^62^ with altered library preparation using the SQKRBK004 rapid barcoding kit. The data was demultiplexed using Deepbinner (v0.2.0)^63^ and basecalled using Guppy (v3.6.0).

For Illumina whole genome sequencing, DNA was extracted from a 10 mL culture using the QIAGEN® Genomic DNA Buffer Set and Genomic-tips™ 100/G set (midi-prep) (QIAGEN, Hilden, Germany). This procedure adhered to the Sample Preparation and Lysis Protocol for Bacteria as outlined in the QIAGEN® Genomic DNA Handbook, with the addition of a preliminary step involving freezing at -20°C. DNA was eluted in 10 mM Tris-HCl (pH 8.5) and stored at -20°C until further processing. Concentrations and quality of the DNA was determined by fluorescence spectroscopy (Qubit™ dsDNA HS assay; Invitrogen by Thermo Fisher Scientific Inc., Eugene, OR, USA) and absorption (DeNovix 439 DS-11+, DeNovix Inc., Wilmington, DE, USA), respectively. The KAPA HYPRplus kit was used to generate illumina libraries which were sequenced at The Novo Nordisk Foundation Center for Biosustainability (Technical University of Denmark, Kgs. Lyngby, Denmark) on the Illumina Miseq 2x300nt PE platform.

The genome assembly was performed by adaptor trimming from the Nanopore data using Porechop (v0.2.4)^64^ and reads smaller than 1,000nt were removed using Filtlong (v0.2.0)^65^ resulting in a total of 1,101,671,292nt in reads with an N50 of 18,196nt, which were assembled de novo with Flye (v2.8-b1674) with the nano-raw setting and 5 polishing iterations^66^resulting in a 13Mb assembly. Illumina reads were trimmed using Trim Galore with Cutadapt (v2.10) ^67^ using setting --length 100 and --quality 20. An alignment of Nanopore and Illumina data was created using Bowtie-2 (v.2.3.4.1)^68^ with an overall alignment rate of 98.55%. The Nanopore-only assembly was polished using Illumina data with Unicycler (v0.4.8)^69^ resulting in a chromosome of 11,468,629bp, a mean Nanopore coverage of 70 and a^42^ benchmarking BUSCO (v4.0.5)^70^ score of 99.7% complete genes with 4 duplicate genes using the actinobactera_class ODB10 database. Taxonomical classification was done using GTDB-Tk [67]

### Metabolomic sample preparation

For metabolomic analysis, the *Streptomyces* spp. were incubated in dark for 7 days at 30°C on 90 mm agar plates containing 20 mL SFM or ISP2 media plus agar. The samples were prepared by taking three plugs of 5.5mm diameter and transferring them to a 2mL Eppendorf tube (VWR Chemicals; 211-2120). The plugs were then submerged in 1mL ethyl acetate (VWR Chemicals; 34858) and exposed to ultrasonication for 60 minutes. The mixture was then transferred to a clean Eppendorf tube (VWR Chemicals; 211-2120) and evaporated under nitrogen. Samples were redissolved into 200µL of methanol (Sigma-Aldrich; 34860) and ultrasonicated for 15 minutes. The mixture was then centrifuged at 14000 RCF for 3 minutes and the supernatant transferred to HPLC vials (Thermo Fisher Scientific; C4000-12) and sealed with a cap (Thermo Fisher Scientific; 9-SCKGST1) and subjected to ultrahigh-performance liquid chromatography-high resolution electrospray ionization mass spectrometry (UHPLC-HRESIMS) analysis.

### Identification and cloning of the putative azodyrecin biosynthetic gene cluster

The putative azodyrecin BGC was identified by alignment of known azoxy producing BGCs, or their whole genome sequences (WGS) to the *S. mirabilis* P8-A2 WGS using LASTZ^71^. The query sequences were: valanimycin BGC (NCBI: AY116644.1), *S. griseofuscus* 7434AN4 WGS (NCBI: GCF_008064995.1) and azodyrecin BGCs from *Streptomyces sp.* A1C6 (NCBI: LC712331) and *Streptomyces sp.* RM72 (NCBI: LC712332). The identified regions were verified using antiSMASH 7.0^43^ that finds *azdB* and classifies cluster in “Other” group. The identified region was compared to other azoxy producers using clinker^46^. The gene functions were predicted using InterPro^49^ scan v. 5.63-95.0 for final insights in candidate cluster and decision on cluster borders.

The putative azodyrecin BGC was cloned according to modified^47^ CATCH-cloning^72^ procedure. To capture the azodyrecin BGC, Cas9 protospacer sequences were designed using Geneious Prime to find sgRNA bunding sites and score them based on their specificity^73^ and activity^74^. The sgRNAs were ordered as 2 nmol Alt-R CRISPR-Cas9 lyophilised and ready-to-dissolve fragments from Integrated DNA Technologies. The sgRNAs contained following protospacer sequences, sgRNA001: “GCGCATCCGTGACACCACCG” and sgRNA002: “ACGCTGGCGTCACCAACGCG”. The primer extension PCR of pXJ157 plasmid was performed using matmal0031 “TTCTTCCAGGAGCACGCTGGCGTCACCAACAATTGTTATCCGCTCACAATTCC” and matmal0032 “CGTTCACCGGCATGCGCATCCGTGACACCACTTTAAGAAGGAGATATAC-CATGAGC” primers. Plasmid assemblies were verified by restriction enzyme mapping using PagI and SgsI (Figure S.6, Figure S.7) and whole plasmid sequencing using Oxford Nanopore Flow Cell (R9.4.1) and mapping to reference plasmid using minimap2^75^. The generated plasmids IDs are pAzd (pXJ157-Azodyrecin_BGC) (Figure S.1) and pAzdΦC31 (pXJ157-*apr*-*attp*-*int*-Azodyrecin_BGC) (Figure S.2).

### Knock-out strain construction

Gene knockouts in *S. mirabilis* P8-A2 were performed by introduction of stop codons near the N-terminus of the target coding sequence using CRISPR-cBEST base editing^40^. The *S. mirabilis* P8-A2 KO strains were generated as previously described^41^ with the following alterations. Instead of pCRISPR-cBEST (RRID: <u>Addgene_125689</u>) we created a new set of plasmids containing dual restriction sites in the protospacer cloning site and with either *erm*EP*, *kasO*P*, or SF14P promoters driving sgRNA transcription. Using NcoI and NheI instead of only NcoI reduces the number of negative colonies and greatly simplifies the cloning of many base editing plasmids. The new plasmids were named pCRISPR-cBESTv2-PermE* (pCW135, Addgene ID: 209446) (Figure S.3), pCRISPR-cBEST-v2-kasOP* (pCW136, Addgene ID: 209447) (Figure S.4) and pCRISPR-cBEST-v2-SF14P (pCW137, Addgene ID: 209448) (Figure S.5). The pCW135 and pCW136 plasmids were constructed by primer extension PCR using original the original pCRISPR-cBEST plasmid as backbone. pCW137 was constructed based on pCW135 using a gBlock from IDT containing the SF14P promoter sequence. All PCRs were performed using Q5 High-Fidelity DNA Polymerase (NEB; M0491). The PCR products were assembled using NEBuilder HiFi DNA Assembly (NEB; E2621) following the suppliers’ specifications. The assembled plasmids were transformed into *E. coli* Mach1 for plasmid propagation. The plasmids were verified by Sanger sequencing (Eurofins Genomics). The backbone for pCW135 and pCW136 was acquired by digestion of pCRISPR-cBEST using BstBI and NcoI. The backbone for construction of pCW137 was obtained through digestion of pCW135 with HindIII and NheI.

The Cas9 sgRNA protospacer sequences were designed using CRISPy-web^76^ . Selection criteria were the absence of 0 bp mismatches, and a location close to the start codon to result in the most truncated protein. A list of protospacer sequences and vector IDs are presented in Table S.4. The oligonucleotides containing 20nt protospacer sequence and 20nt overlaps to the backbone in either side and verification primers were ordered from Integrated DNA Technologies. The assembled CRISPR-cBEST plasmids were verified by Sanger sequencing (Eurofins Genomics) of colony PCR products using primers matmal0137 (TGTGTGGAATTGTGAGCGGATA) and matmal0138 (CCCATTCAA-GAACAGCAAGCAG). Verification of introduced stop codons in *S. mirabilis* P8-A2 exconjugants that showed resistance to apramycin were performed by PCR amplification of the target site and subsequent Sanger sequencing using verification primers presented in Table S.5. After the removal of CRISPR-cBEST plasmids from the strains, the PCR and subsequent Sanger sequencing (Eurofins Genomics) was repeated to ensure maintenance of the installed mutations.

### Phylogenetic distribution of azoxy BGCs

1528 genomic sequences of *Streptomyces* spp. were acquired form NCBI on 16.02.2023, which were selected with requirement to contain up to 100 contigs. Additionally, 30 different genomes from *Actinomycetia* class not belonging to *Streptomycetales* were included to be used as outgroup for rooting of the phylogenetic tree. The resulting dataset had average N50 of 4.5 Mb, and the lowest quality genome had N50 of 109 Kb. We used BGCflow^77^ to systematically acquire nucleotide fasta files from NCBI using ncbi-genome-download (v.0.3.1)^78^ followed by annotation using prokka (v.1.14.5)^79^, with database of HQ manually annotated genomes as described in Gren et al. 2020^80^, including additionally *Streptomyces lividans* TK24 (GCA_000739105.1).

The phylogenetic tree was inferred using autoMLST^57^ as implemented in the autoMLST wrapper^81^. Diamond database was created using cblaster ^82^. The resulting phylogenetic tree was uploaded to iTOL^83^.

To map the clusters onto the phylogenetic tree we wrote a Jupyter notebook, GeneClusterPhyloMapper, that processes input files through clinker^46^ and cblaster^82^ and formats the data for visualization with iTOL^83^. The notebook and documentation is available at: https://github.com/MatissMaleckis/GeneClusterPhyloMapper.

`GeneClusterPhyloMapper` is implemented using python 3.10.9 Jupiter notebook. The notebook requires installation of `os`, `pandas`, `numpy` and `Bio` packages. Within the notebook bash scrips of clinker (v.0.0.28)^46^ and cblaster (v.1.3.18)^82^ are executed, thus environments must be set up prior use. The notebook starts the analysis of user provided input files to find protein similarities between the input BGCs. The notebook compares provided BGC GenBank files using clinker^46^ and shared proteins are grouped. The user is asked to define required number of proteins in a group for BGC to be considered as core (usually same number as input BGCs, thus proteins shared across all BGCs are core). The protein group list is then populated with experimentally validated proteins involved in biosynthesis and not already detected by clinker. The notebook creates input file for cblaster^82^ analysis by extracting protein sequences from the input GenBank files and merge them into single file. The file is then processed by cblaster analysis in local mode using a previously created diamond database. For each cblaster output genome, the notebook calculates how many core protein groups are identified and how many proteins belong to specific BGC. The notebook is then filtered to contain user specified minimum number of core proteins and total proteins detected. Finally, the BGC is assigned based on the ratio between proteins detected belonging to specific BGC and number of BGC specific proteins in the protein group list. If any of the genome has more than one hit, only best hit is kept for further analysis, while excluded hits are saved in separate file for later inspection. Finally, the data is used to generate tables that can be directly imported into iTOL annotation editor. The user has to provide color for each of the input BGC, whereafter notebook creates a table that can be imported in iTOL as color gradient. The notebook also generates a table for shape plot mapping in iTOL. This data maps for each genome the presence absence matrix of each protein group. The data is color coded based on number of records in specific group, while the size of nodes in iTOL represents highest protein identity hit within the group. The generated tables can directly be used in iTOL annotation editor v1.8 for Excel to annotate the phylogenetic tree.

In this study we applied `GeneClusterPhyloMapper` notebook onto valanimycin (NCBI: AY116644.1), azodyrecin (MiBIG: BGC0002805), proposed KA57A BGC (NCBI: GCF_008064995.1 [NZ_AP018517.1 [2,049,086:2,106,642]]) and azoxymycin BGC of *S. chattanoogensis* NRRL ISP-5002 (NCBI: GCF_001294335.1 [NZ_LGKG01000136 [21,828:35,070]]). We provided a list of experimentally validated protein sequences involved in azoxy compound biosynthesis (NCBI: AN10236.1, AAN10237.1, AAN10239.1, AAN10241.1, AAN10242.1, AAN10243.1, AAN10244.1, AAN10246.1, AAN10247.1, AAN10248.1, AAN10249.1, PU648_09290, PU648_09235, PU648_09240, PU648_09245, PU648_09255, PU648_09260, PU648_09270, PU648_09275, PU648_09280, PU648_09285, PU648_09325, PU648_09360, KP687735.1, KP687738.1, KP687739.1, KP687742.1). We defined cluster detection requirements as follows: `proteins shared to be core` = 3, `minimum core proteins` = 4

## Data availability statement

The data underlying this study is openly available. The genomic data has been deposited in the NCBI and MiBIG database under accession number JARAKF000000000 and BGC0002805 respectively. The metabolomic data has been deposited in MassIVE under ID MSV000092718. The notebook for mapping of clusters onto iTOL phylogenetic tree is available on GitHub [https://github.com/MatissMaleckis/GeneClusterPhyloMapper]. The iTOL phylogenetic tree (named: Azoxy-like BGC Distribution) can be explored under user ID matmal [https://itol.embl.de/shared/matmal].

## Acknowledgment

This study was supported by the Danish National Research Foundation (DNRF137) as part of the Center for Microbial Secondary Metabolites (CeMiSt). T.W. would furthermore acknowledge funding by the Novo Nordisk Foundation (NNF20CC0035580, NNF16OC0021746).

The metabolomic data was generated at DTU Metabolomics Core facilities with help of Aaron J. C. Andersen. For the technical advice with metabolomic analysis we would like to thank Eftychia E. Kontou, for sequencing assistance Alexandra Hoffmeyer and Oliwia Vuksanovic, for assistance with genome mining and data treatment Matin Nuhamunada, Omkar Mohite, Simon Shaw and Kai Blin and with genome engineering of *Streptomyces* spp. Renata Sigrist, Zhijie Yang, Subhasish Saha and Peter Gockel.

