## Supplementary Information for "Biosynthesis of the azoxy compound azodyrecin from *Streptomyces mirabilis* P8-A2"

### Azoxy producers in *Streptomyces* spp. overview

Table S.1**:** Azoxy compounds isolated from *Streptomyces* spp., their sequence availability and the genomic sequence that was used in this study.

| Azoxy compou | Producing microorgan-ism | Sequence availability: | Nucleotide se-quence used: | |
|---|---|---|---|---|
| Azodyrecins A-C | *S. mirabilis* P8-A2 | WGS | This Paper | 18 |
| Azodyrecins A-F | *Streptomyces* sp. A1C6, *Streptomyces* sp. RM72 | BGC | LC712331 LC712332 | 19 |
| Valanimycin | *S. viridifaciens* MG456-hF10 | BGC | AY116644.1 | 34 |
| KA57A | (*) *S. rochei* 7434AN4 | WGS | GCF_008064995.1 | 13,45 |
| Unnamed | *S. avermitilis* MA-4680 | WGS | GCF_000009765.2 | 24 |
| Azoxymycins | *S. chattanoogensis* L10 | Genes only (Table S.2) | *S. chattanoogensis* NRRL ISP-5002 GCF_001294335.1 | 20 |
| Elaiomycin | *S. gelaticus* C2828-PD-04942 | n.a. | n.a. | 7–9 |
| Elaiomycins B and C | *Streptomyces* sp. BK 190 | 16S rRNA: *S. atratus* NRRL B-16927 | n.a. | 3,4 |
| Elaiomycins D–F | *Streptomyces* sp. HKI0708 | | n.a. | 5 |
| Elaiomycins K and L | *Streptomyces* sp. Tü 6399 | | n.a. | 6 |
| Maniwamycins A-B | *S. prasinopilosus* KC-7367 | n.a. | n.a. | 15 |
| Maniwamycins A-G | *Streptomyces* sp. TOHO-M025 | 16S rRNA: (**) *S. hygro-scopicus* subsp. TL01 and 5008 | n.a. | 14,16 |
| O-Alkylazoxymy-cins A–F | *Streptomyces* sp. Py50 | 16S rRNA: *S. atratus* NBRC 3897 | n.a. | 21 |
| Geralcin C | *Streptomyces* sp. LMA-545 | 16sRNA: Low quality | n.a. | 23 |
| Jietacins | *Streptomyces* sp. KP-197 | n.a. | n.a. | 17 |
| LL-BH872α | *S. hinnulinusn* n.s. [Lederle Culture No. BH872] | n.a. | n.a. | 10 |
| DC-8118 A and B | *Streptomyces* sp. DO-118 | n.a. | n.a. | 22 |

[*] Organism misclassified: declared type ANI 85.39% (*S. rochei*), best match type ANI 99.64% (*S. griseofuscus*).
[**] Organisms misclassified: declared type ANI 84.02% (*Streptomyces hygroscopicus subsp. hygroscopicus*), best match type ANI 98.96% (*S. corchorusii*).

Table S.2 Published azoxymycin biosynthetic gene similarity to homologue genes identified within 14kb region in the *S. chattanoogensis* NRRL ISP-5002, GCF_001294335.1 [NZ_LGKG01000136 [21,828:35,070]].

| Azoxymycin BGC gene | Pairwise nucleotide identity (%) | Pairwise protein identity (%) |
|---|---|---|
| *azoA* | 97.9 | 96.2 |
| *azoB* | 98.5 | 99.2 |
| *azoC* | 98.5 | 98.8 |
| *azoD* | 98.6 | 98.7 |
| *azoE* | 98.5 | 99.4 |
| *azoF* | 98.9 | 98.3 |
| *azoG* | 98.8 | 99.2 |
| *azoH* | 99.6 | 100.0 |
| *azoI* | 99.3 | 98.9 |
| *azoJ* | 98.4 | 98.9 |
| *azoK* | 99.1 | 99.1 |
| *azoL* | 98.5 | 96.5 |
| *azoM* | 98.9 | 98.7 |
| *azoN* | 98.8 | 97.9 |
| *azoO* | 99.3 | 99.1 |

Table S.3: Overview characterized gene functions from valanimycin biosynthesis studies and the genes shared with azodyrecin producer *S. mirabilis* P8-A2 and KA57-A producer *S. griseofuscus* 7434AN4.

| **Valanimycin genes** | **Azodyrecin genes** | **KA57-A locus tags** | **Identified function in valanimycin biosynthesis** | |
|---|---|---|---|---|
| *vlmA* | *azdA* | PBHDNOKL_17950 | AA-tRNA-dependent transferase | [34,36] |
| *vlmB* | *azdB* | PBHDNOKL_17840 | - | - |
| *vlmH* | *azdH* | PBHDNOKL_17790 | Isobutylamine N-hydroxylase | [28,34] |
| *vlmO* | *azdO* | PBHDNOKL_17820 | - | - |
| *vlmR* | *azdR* | PBHDNOKL_17830 | FAD reductase | [27,34] |
| *vlmC* | - | PBHDNOKL_17880 | Amino acid permease | [34] |
| *vlmD* | - | - | Valine decarboxylase | [34] |
| *vlmE* | - | - | *tetR* regulator | [34] |
| *vlmF* | - | - | Valanimycin resistance - DHA12 transport protein | [30,34] |
| *vlmG* | - | - | - | - |
| *vlmI* | - | PBHDNOKL_17960 | SARP-regulator | [29,34] |
| *vlmJ* | - | PBHDNOKL_17680 | diacylglycerol kinases | [34,37] |
| *vlmK* | - | PBHDNOKL_17660 | MmgE/PrpD superfamily of proteins | [34,37] |
| *vlmL* | - | PBHDNOKL_17890 | Seryl tRNA synthetase | [34,35] |

### Plasmid maps

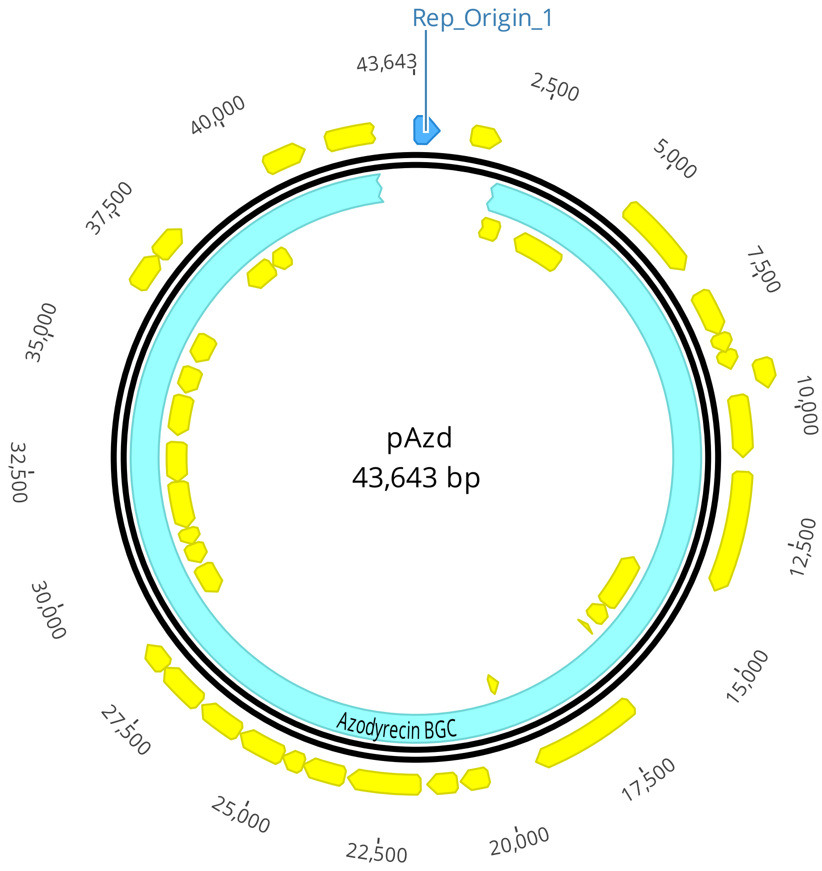

Figure S.1: Plasmid map of pAzd (pXJ157-Azodyrecin_BGC). Generated using Geneious version 2023.2 created by Biomatters.

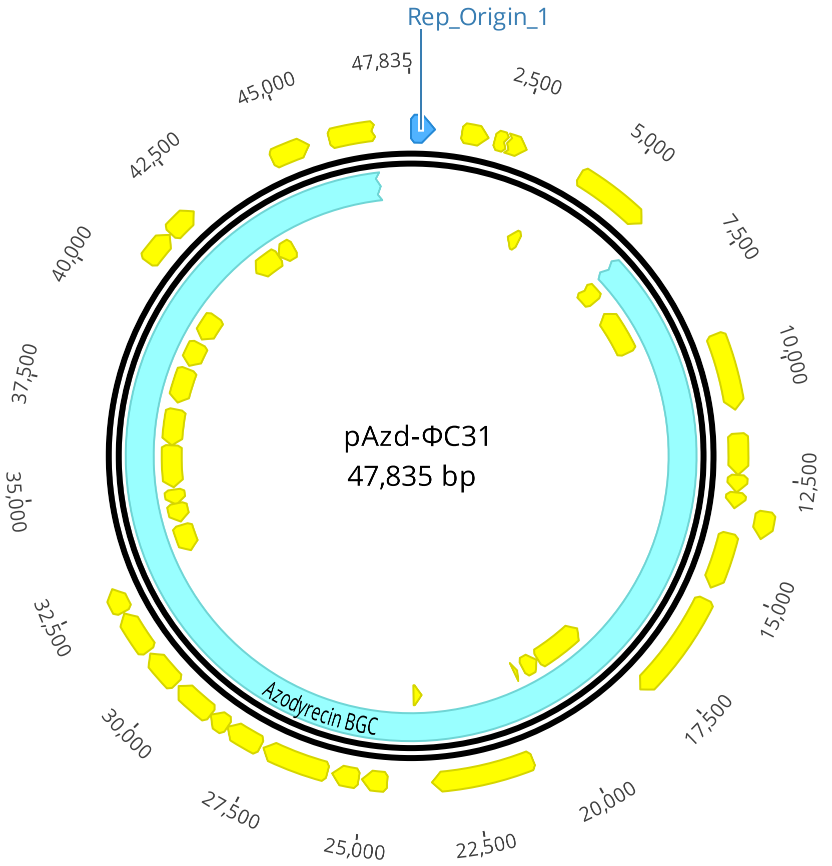

Figure S.2: Plasmid map of pAzd-ΦC31(pXJ157-*apr*-*attp*-*int*-Azodyrecin_BGC). Generated using Geneious version 2023.2 created by Biomatters.

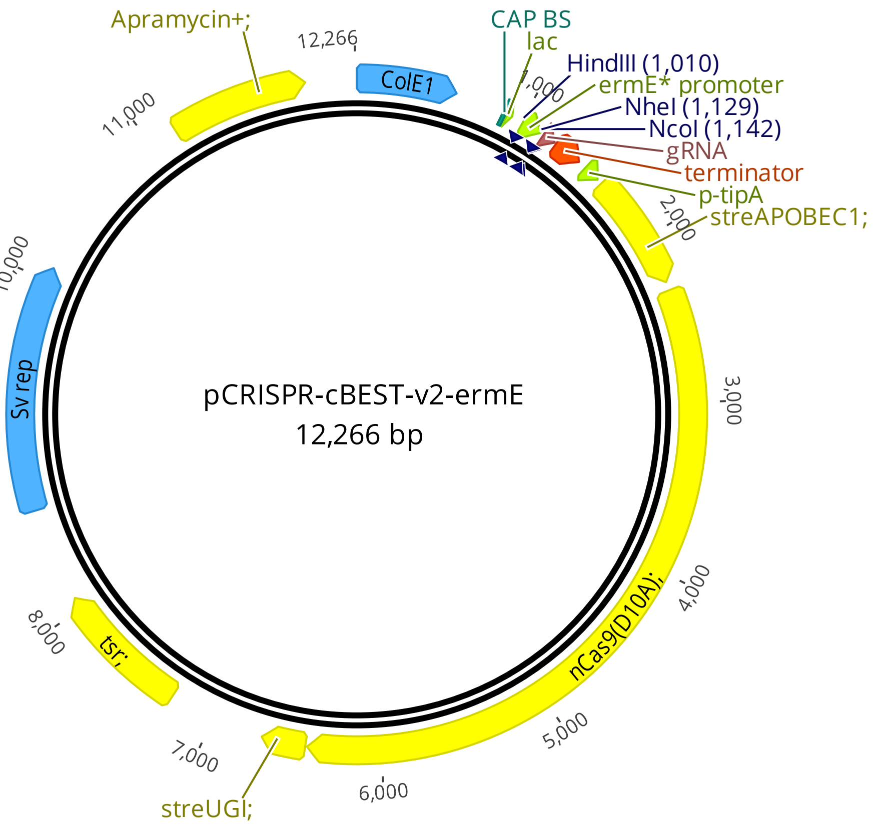

Figure S.3: Plasmid map of pCRISPR-cBEST-v2-ermE (pCW135). Generated using Geneious version 2023.2 created by Biomatters.

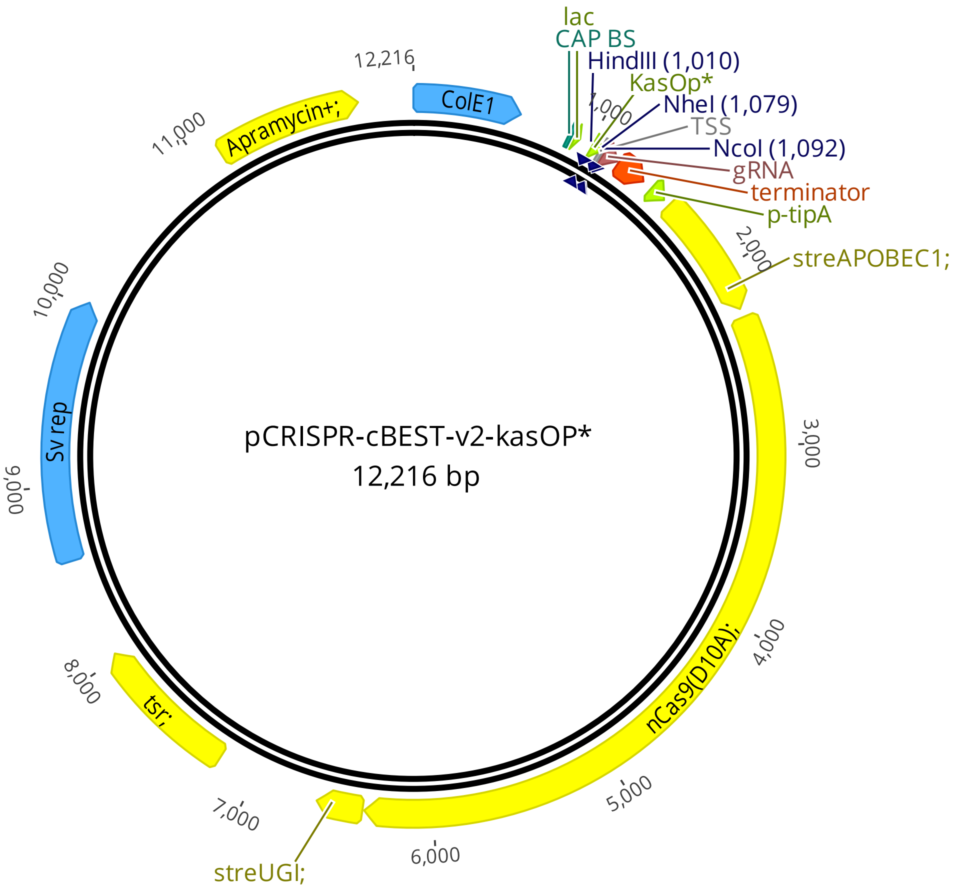

Figure S.4: Plasmid map of pCRISPR-cBEST-v2-kasOP* (pCW136). Generated using Geneious version 2023.2 created by Biomatters.

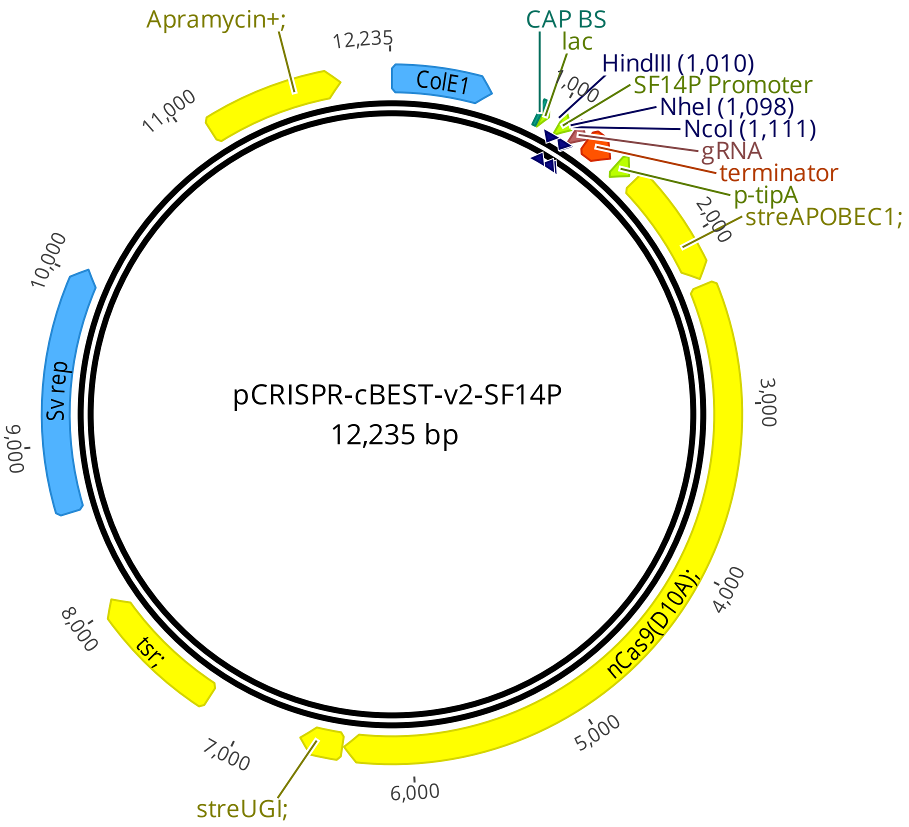

Figure S.5: Plasmid map of pCRISPR-cBEST-v2-SF14P (pCW137). Generated using Geneious version 2023.2 created by Biomatters.

### Heterologous expression of the cluster

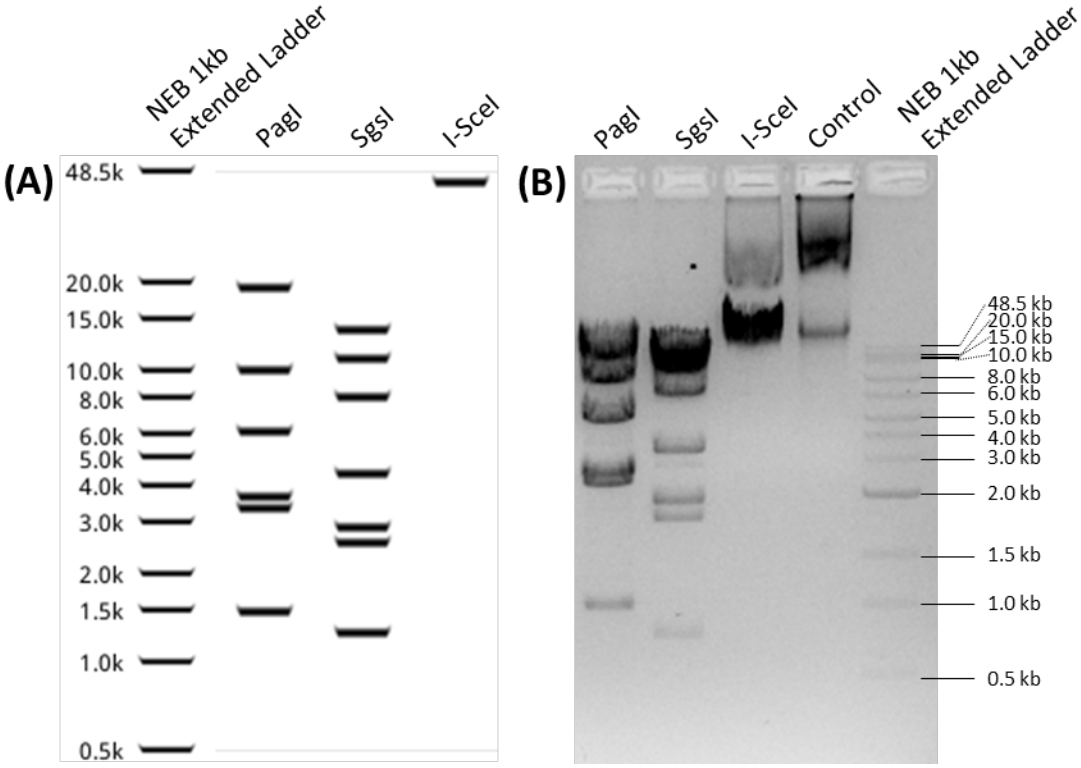

Figure S.6: Restriction map of pAzd plasmid containing Azodyrecin BGC. **(A)** is expected virtual gel that is generated using Geneious Prime software, showing the cleavage using eater PagI, SgsI or I-SceI restriction enzymes that cleave 6, 7 and 1 time respectively and the NEB 1kb Extended Ladder to asses the fragment sizes. **(B)** shows the acquired fragmentation of the digested plasmids using previously mentioned enzymes, with additional loading of control which refers to undigested plasmid.

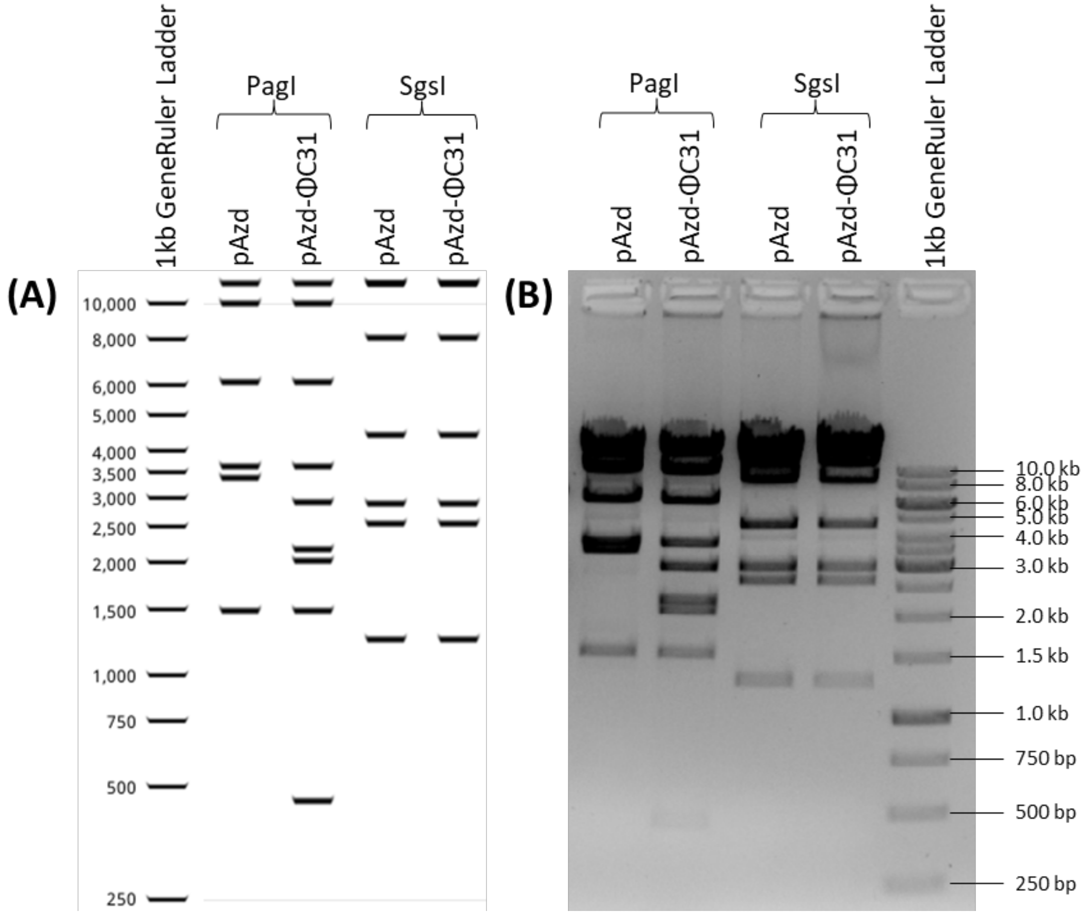

Figure S.7: Restriction map of pAzd and pAzd-ΦC31 plasmid containing Azodyrecin BGC and for latter plasmid also Streptomyces integrating cassette. **(A)** is expected virtual gel that is generated using Geneious Prime software, showing the cleavage using eater PagI and SgsI restriction enzymes and the NEB 1kb Extended Ladder to assess the fragment sizes. **(B)** shows the acquired fragmentation of the digested plasmids using previously mentioned enzymes, with additional loading of control which refers to undigested plasmid.

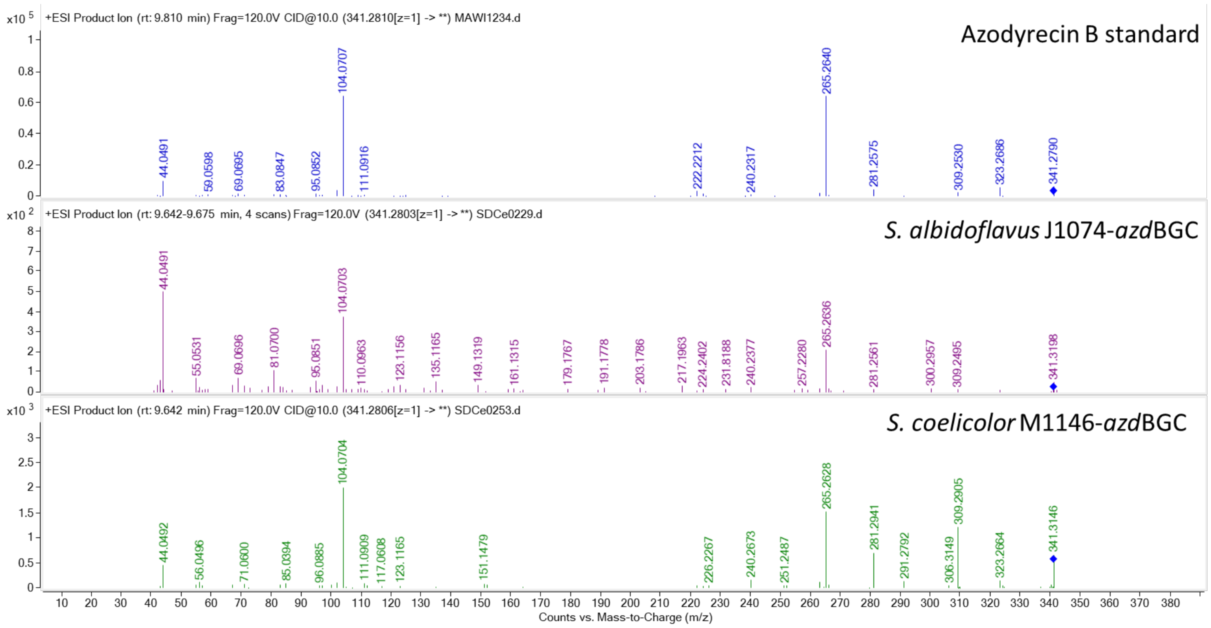

Figure S.8: MS/MS spectra of azodyrecin B (m/z = 341.281) in *S. albidoflavus* J1074 and *S. coelicolor* M1146 vs standard.

### Azodyrecin knock out studies

Table S.4: List of sgRNA protospacer sequences, the CRISPR-cBEST vector plasmid used and the crispy-web predicted specificity/mismatch of the protospacer.

| | | | | Mismatches | | |
|---|---|---|---|---|---|---|
| **Gene target** | **sgRNA Protospacer sequence** | **Vector plasmid** | **PAM** | **0bp** | **1bp** | **2bp** |
| azdA | CCGACGACACCTGTTCCAAC | CW-137 | TGG | 0 | 2 | 45 |
| azdB | GCTGCGCCAGGCCGTCATGC | CW-135 | AGG | 0 | 8 | 195 |
| azdH | GAGAACCAGCTGATGGCGTC | CW-136 | CGG | 0 | 3 | 75 |
| azdN | CATGAACCAGCGGTTGCTAC | CW-135 | GGG | 0 | 1 | 36 |
| azdO | GGCGACCCACCAGCGCACCT | CW-135 | CGG | 0 | 4 | 75 |
| azdP | TGACCCAGGCGGACTTCACC | CW-135 | CGG | 0 | 16 | 215 |
| azdQ | GGTGAACACCCACGCGCCGT | CW-135 | CGG | 0 | 7 | 100 |
| azdR | CAGGGTGTCCCAGCCGGCGG | CW-135 | CGG | 0 | 24 | 293 |
| azdS | AGACCCCAGTCGTCCGTCTC | CW-135 | GGG | 0 | 4 | 63 |
| azdT | CACCTTCCACCAGGTGCCGC | CW-135 | CGG | 0 | 15 | 244 |
| azdU | GAGACGAATCCCCCGTCAGG | CW-135 | CGG | 0 | 1 | 56 |
| azdV | ATCGAGCAGTTGTACGTCAT | CW-135 | CGG | 0 | 3 | 24 |
| azdW | GGTGAACCAGGCGATGTCCG | CW-135 | TGG | 0 | 9 | 199 |
| PU648_09335 | GCTCCAACTGCTGCAGAACG | CW-135 | CGG | 0 | 19 | 153 |

Table S.5: Overview of constructed mutant strains using CRISPR-BEST base editing, the targeted gene in the strain, resulting protein level mutations (* indicates STOP codon) in that gene and the mutation verification primers and their sequences.

| Strain ID: | Gene locus tag and assigned name | Protein level modification | Gene mutation verification primer set and their sequences |
|---|---|---|---|
| SMCe01279 | PU648_09225 (*azdN*) | W58* | matmal0210: CGAACTCCCAGATGGATATGT<br>matmal0211: GTCGTATTCACCGGTCAAC |
| SMCe01277 | PU648_09235 (*azdH*) | Q166* | matmal0103: AAGGAGCAGGGGACGACA<br>matmal0104: CAGCAGCTCGAACCAGACA |
| SMCe01280 | PU648_09240 (azdP) | T75I, Q76* | matmal0212: GTCGAACAACAAGAACCGAG<br>matmal0213: GGTGCTCATGTGGAAGAAC |
| SMCe01281 | PU648_09245 (azdQ) | W74* | matmal0214: GAACTGCTGTTGGTCTTCC<br>matmal0215: GAAGGGCTGTTGGTACTTG |
| SMCe01282 | PU648_09255 (azdR) | W21*, D22N | matmal0216: CTACCAGCGAATCCGTGATTTC<br>matmal0217: TCTCTGATGCTCCGTCAAC |
| SMCe01283 | PU648_09260 (azdS) | W49* | matmal0218: GTTGACGGAGCATCAGAGA<br>matmal0219: GAGGTACTTGAGGTAGTAGTGC |
| SMCe01284 | PU648_09270 (azdT) | W67* | matmal0220: TAGAAGAAGTCGCCGATGTAT<br>matmal0221: TCAACTCCTCACCGTATGAA |
| SMCe01275 | PU648_09275 (azdB) | R72C, Q73* | matmal0079: CTTCACCCCGTTCGTCCA<br>matmal0080: TAGTCGCGTTCCTTCTCCA |
| SMCe01276 | PU648_09280 (azdA) | R39* | matmal0087: AGGCGCCGAACTGGTTGA<br>matmal0088: TGCGCCTCGAATACCTGCT |
| SMCe01274 | PU648_09285 (azdO) | W41*, V42I | matmal0087: AGGCGCCGAACTGGTTGA<br>matmal0088: TGCGCCTCGAATACCTGCT |
| SMCe01278 | PU648_09315 (azdU) | R3* | matmal0226: TTGTTGAGGCTGTAGGTCTT<br>matmal0227: GAATGACATGTCTGGGTCAAG |
| SMCe01285 | PU648_09335 | Q69* | matmal0228: GCGATGAGGTAATTGGAGAC<br>matmal0229: CTATTACGAGTACATGCGTCAC |
| SMCe01286 | PU648_09360 (azdV) | Q109* | matmal0230: GTAGATCTGGTTGAACTCGAAC<br>matmal0231: CAACATCGGCATCAACTACAT |
| SMCe01287 | PU648_09365 (azdW) | W60* | matmal0232: CAGCTCATTGATTGTACCGAC<br>matmal0233: GGTACTGATGGCCATGAAC |

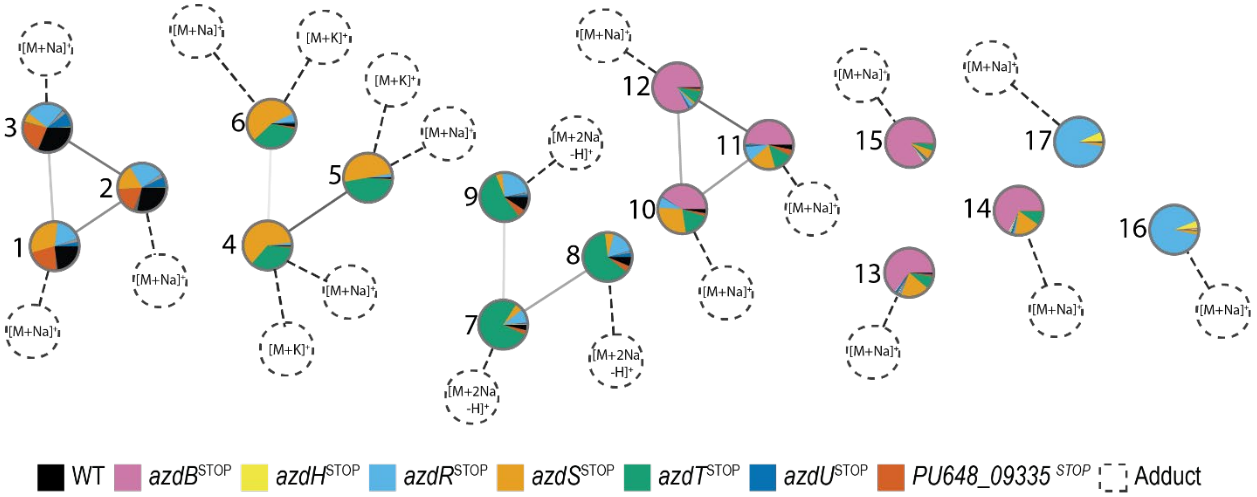

Figure S.9: GNPS clustering of Azodyrecins A-F (**1**-**6**) and the proposed structures of **7**-**17**. Nodes with PI-chart indicate relative amounts of compound detected in specific strains of *S. mirabilis* P8-A2. The different adducts detected are visualized with dotted connections and dotted circles.

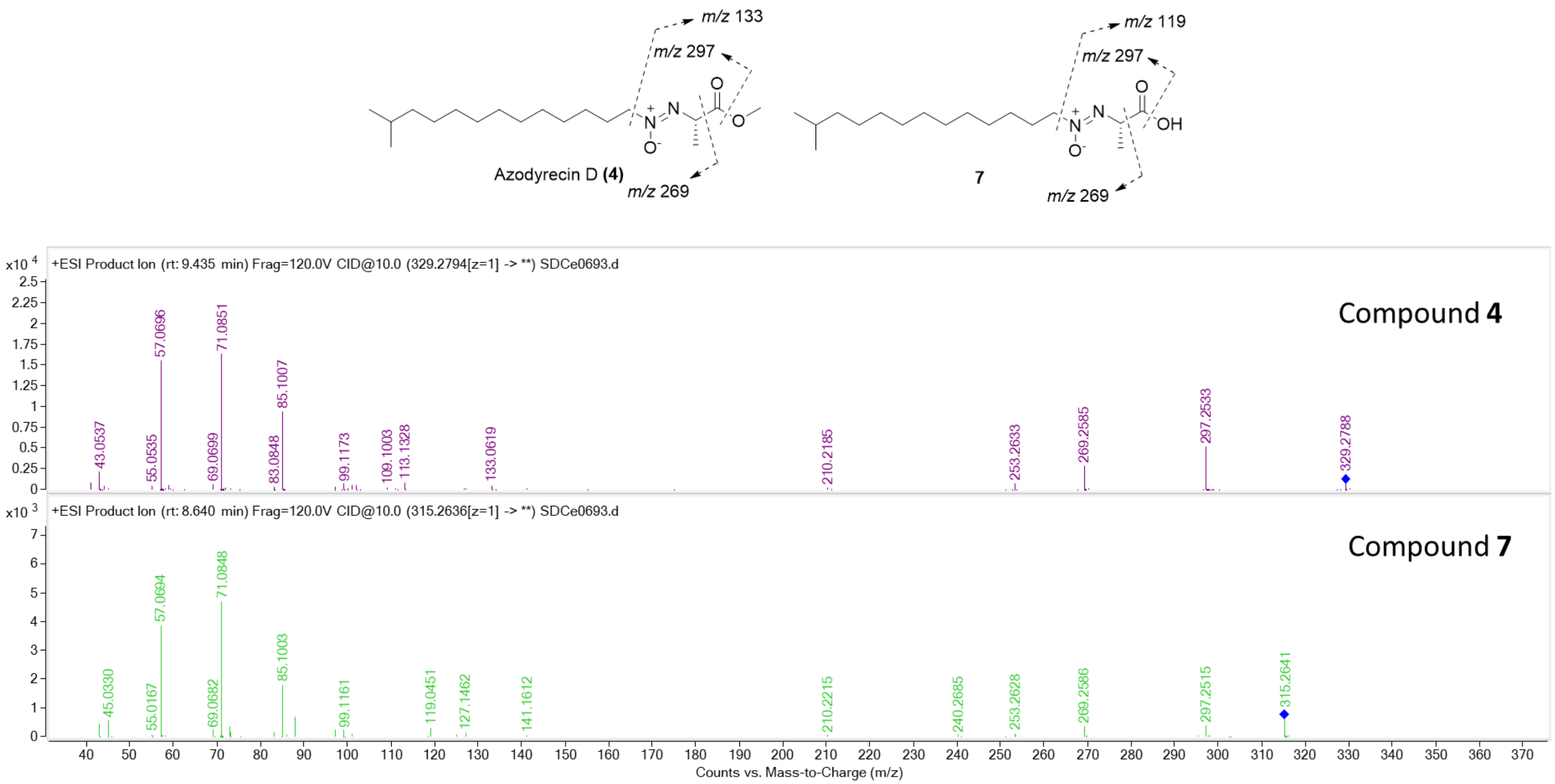

Figure S.10: MS/MS fragmentation spectra comparison of compound 4 - 7.

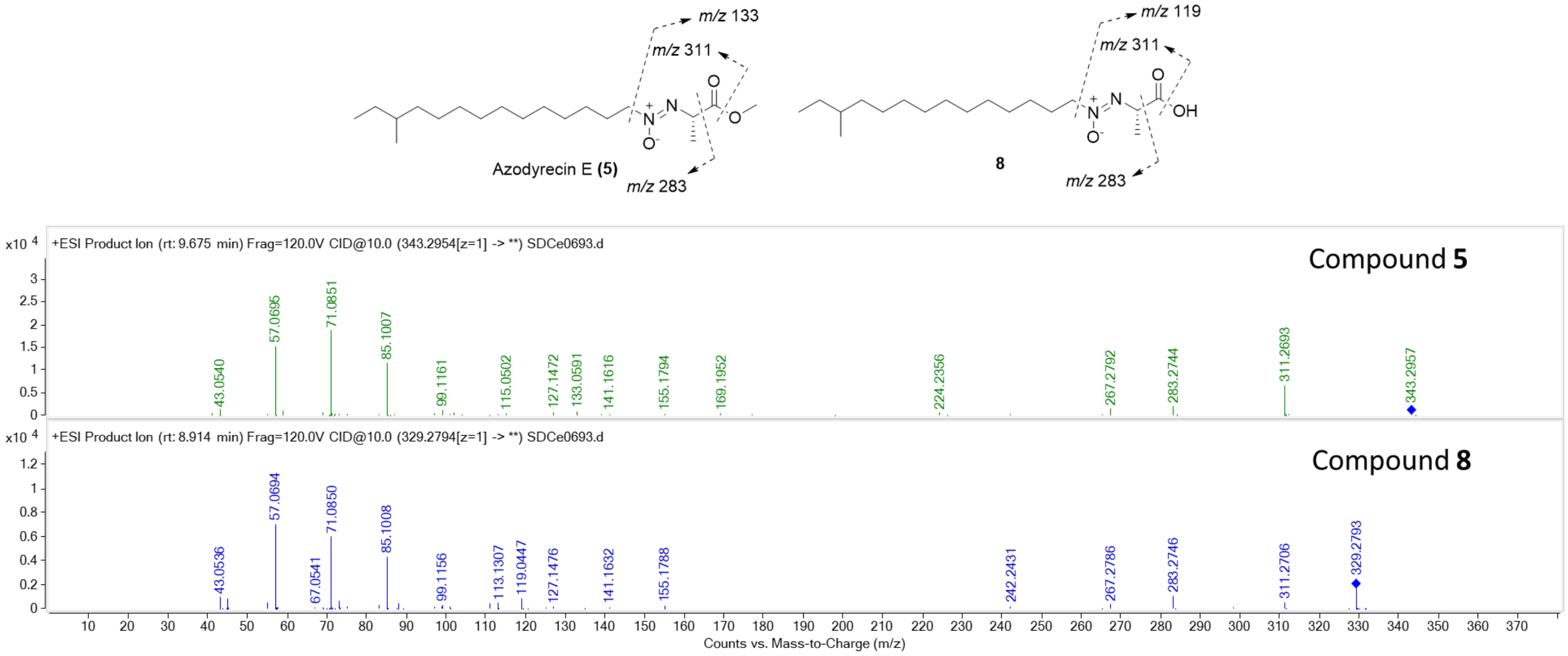

Figure S.11: MS/MS fragmentation spectra comparison of compound 5 - 8.

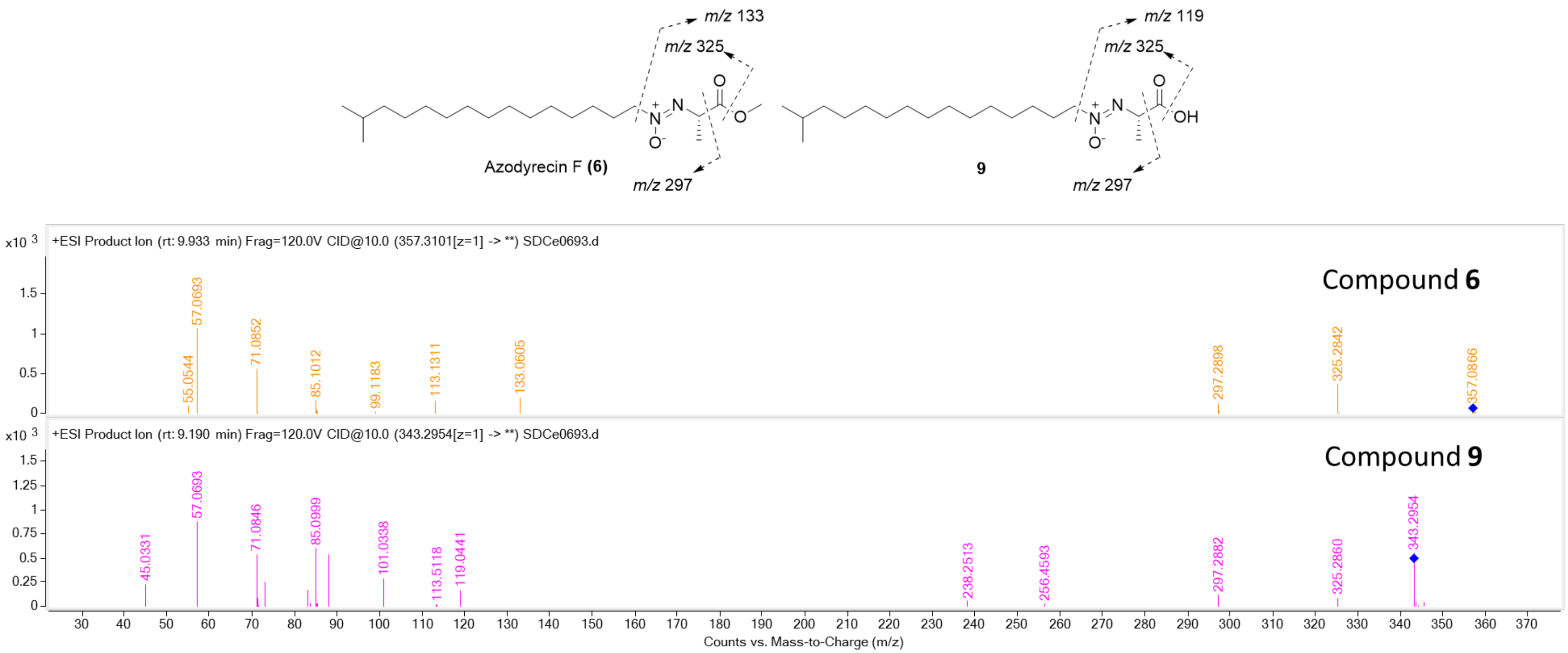

Figure S.12: MS/MS fragmentation spectra comparison of compound 6 - 9.

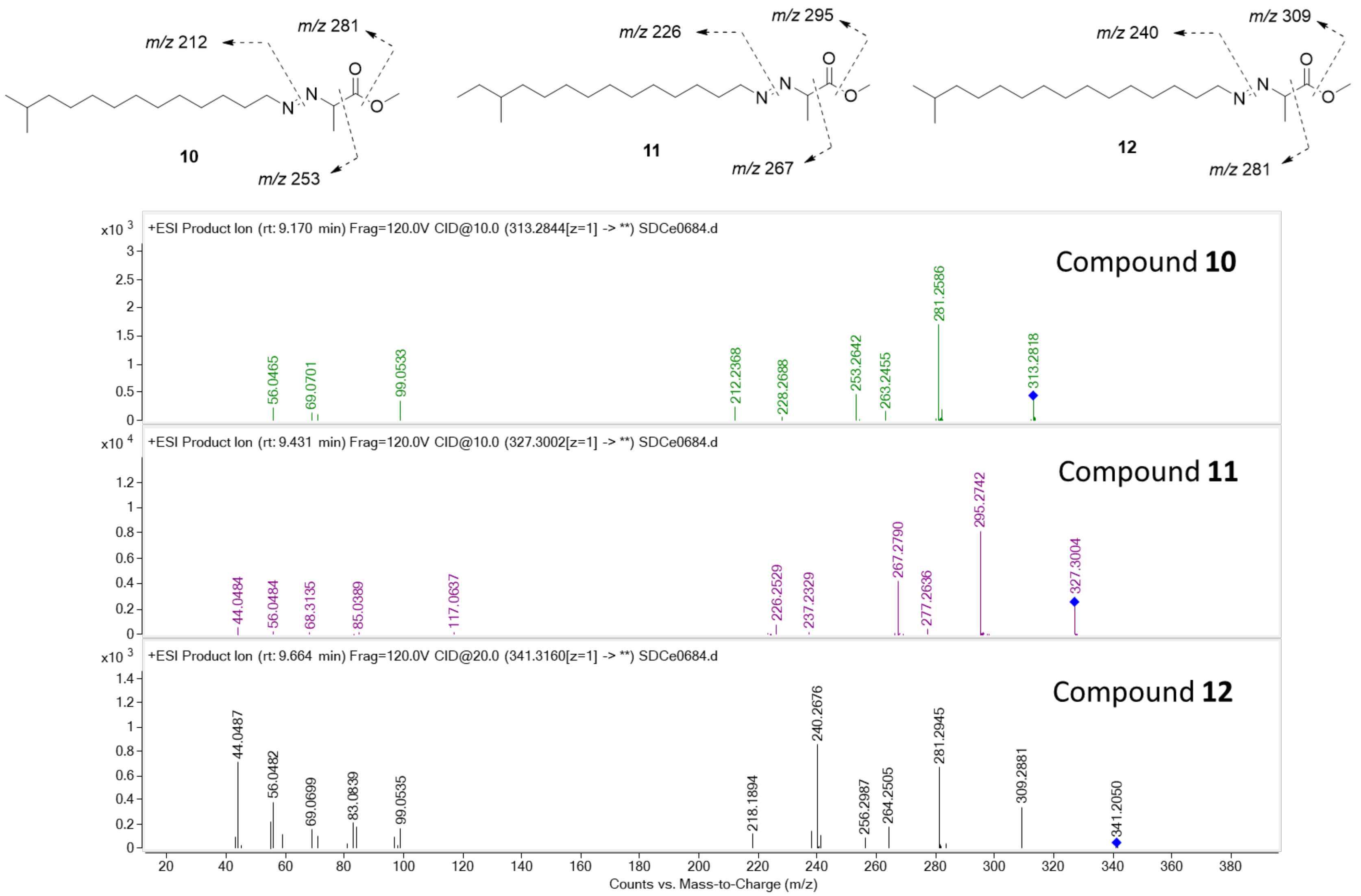

Figure S.13: MS/MS fragmentation spectra comparison of compound 10 - 12, note the shift of 14 masses (CH2) between the compound spectra.

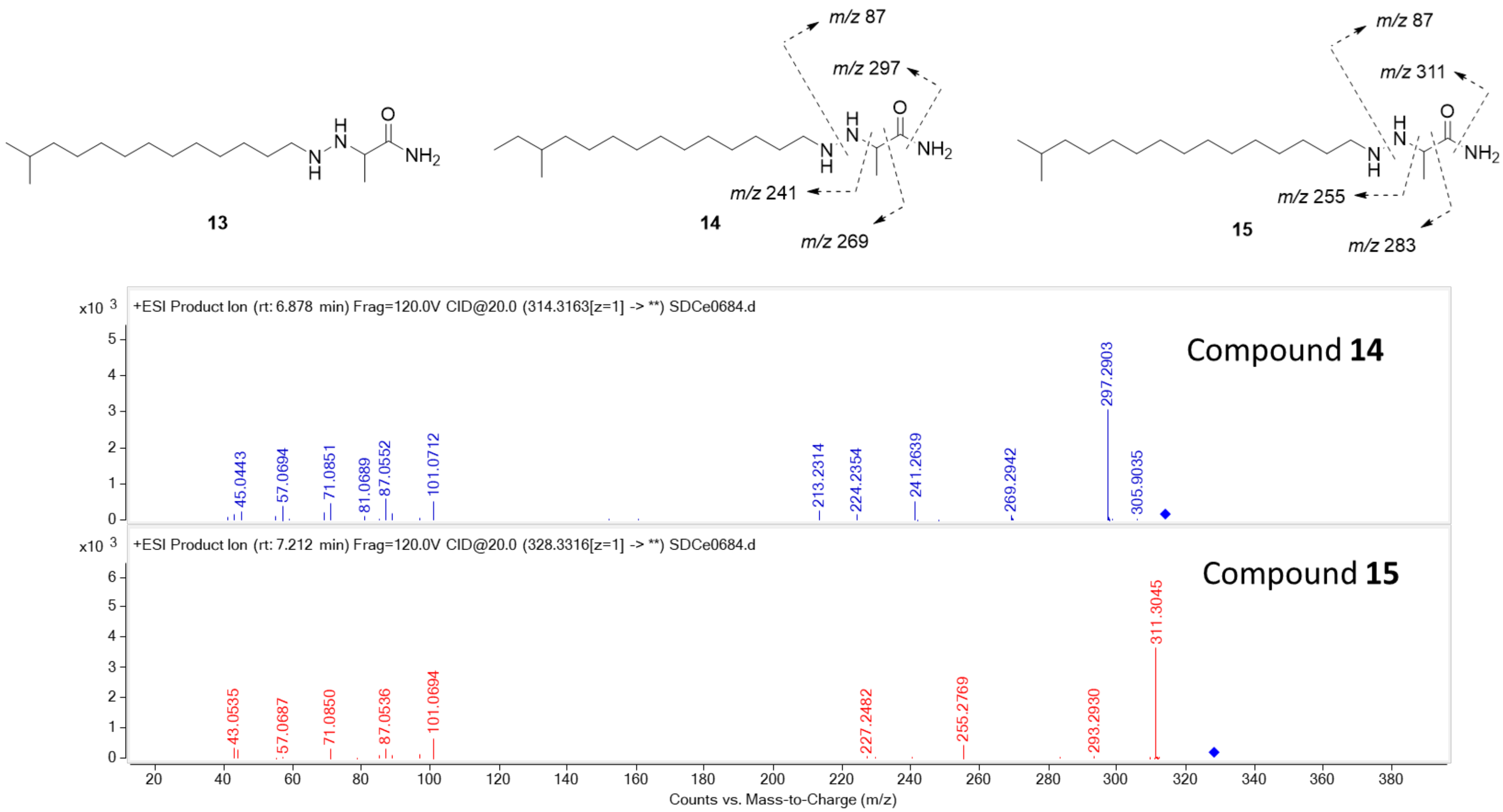

Figure S.14: MS/MS fragmentation spectra comparison of compound 14 and 15, note the shift of 14 masses (CH2) between the compound spectra. Instrument did not collect fragmentation data for compound 13, while MS data was detected.

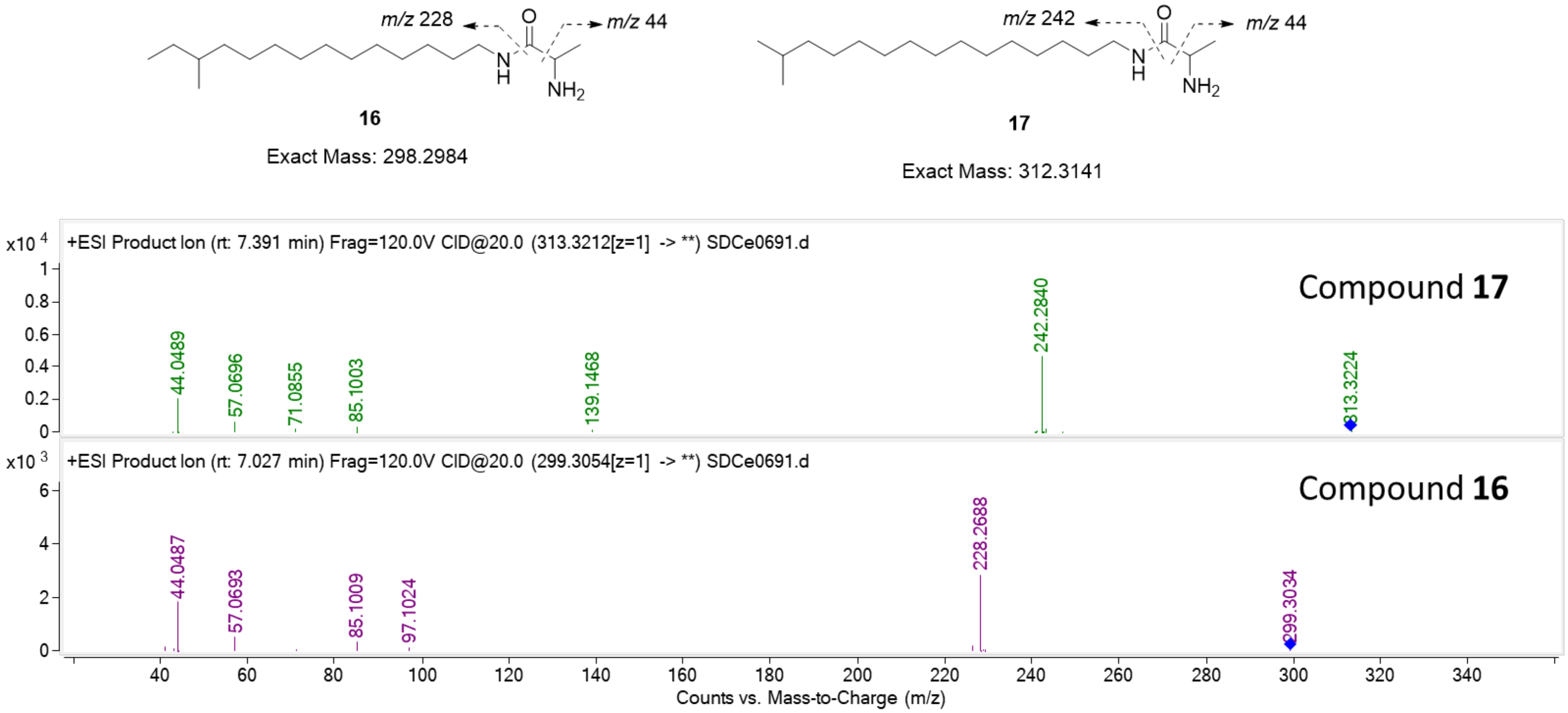

Figure S.15: MS/MS fragmentation spectra comparison of compound 16 and 17, note the shift of 14 masses (CH2) between the compound spectra.

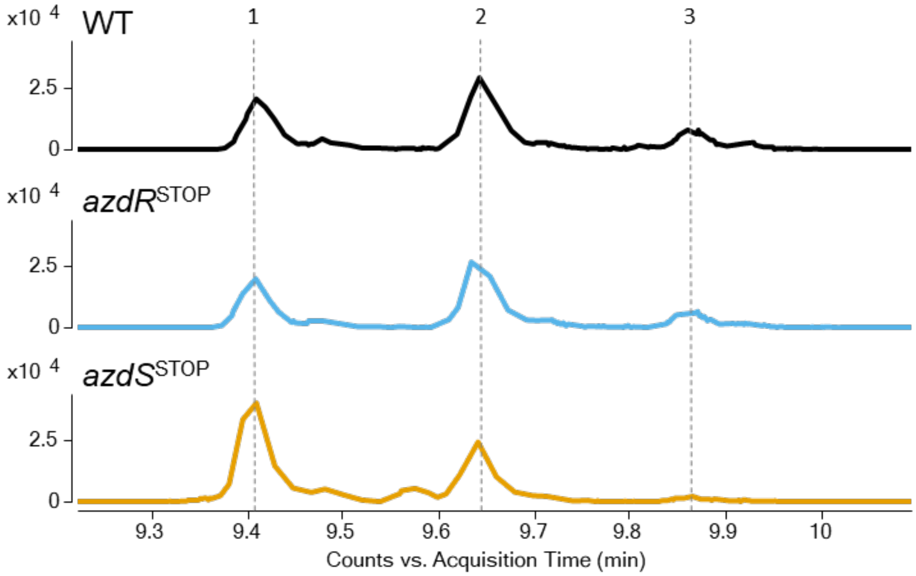

Figure S.16: Extracted ion chromatogram of **azodyrecins A-C (1-3)**, where the first and the last eluting peak correspond to the smallest and the largest EIC respectively.

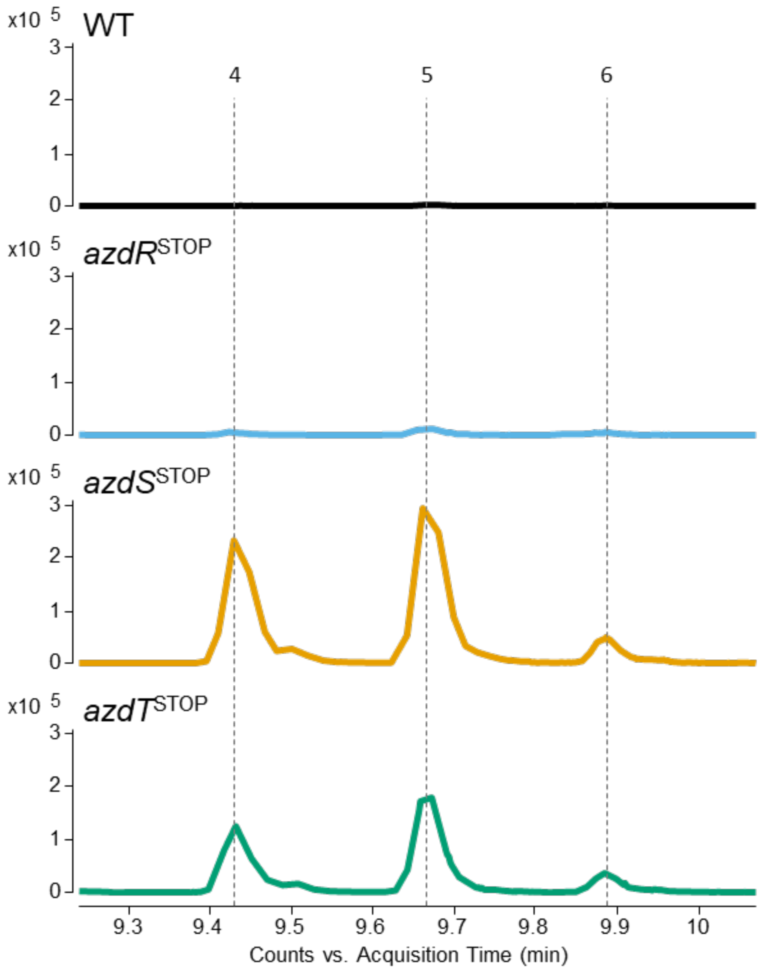

Figure S.17: Extracted ion chromatogram of **azodyrecins D-F (4-6),** where the first and the last eluting peak correspond to the smallest and the largest EIC respectively.

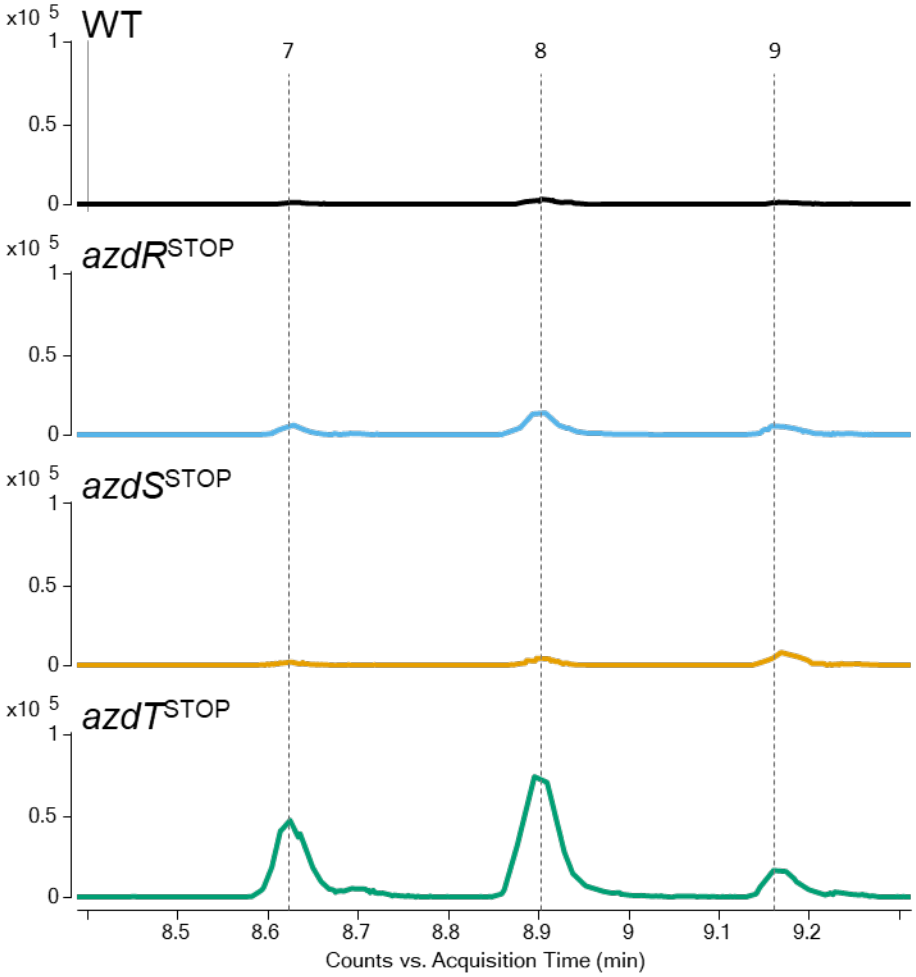

Figure S.18: Extracted ion chromatogram of compound **7-9**, where the first and the last eluting peak correspond to the smallest and the largest EIC respectively.

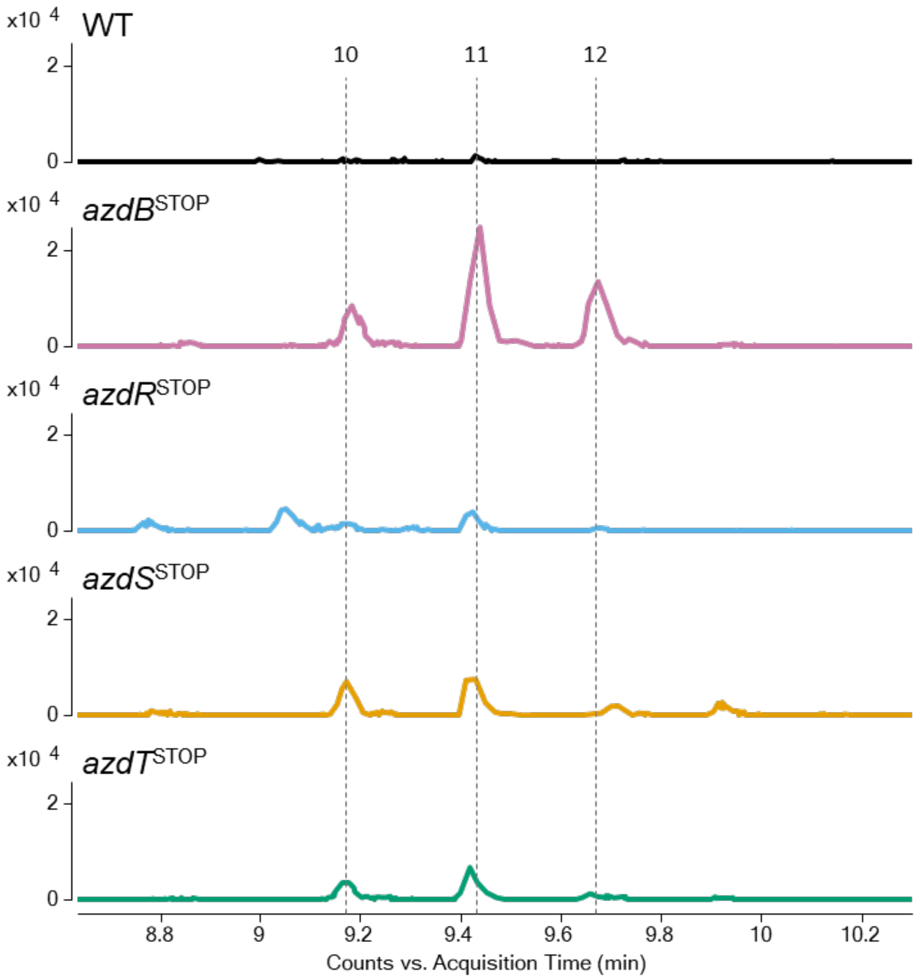

Figure S.19: Extracted ion chromatogram of compound **10-12**, where the first and the last eluting peak correspond to the smallest and the largest EIC respectively.

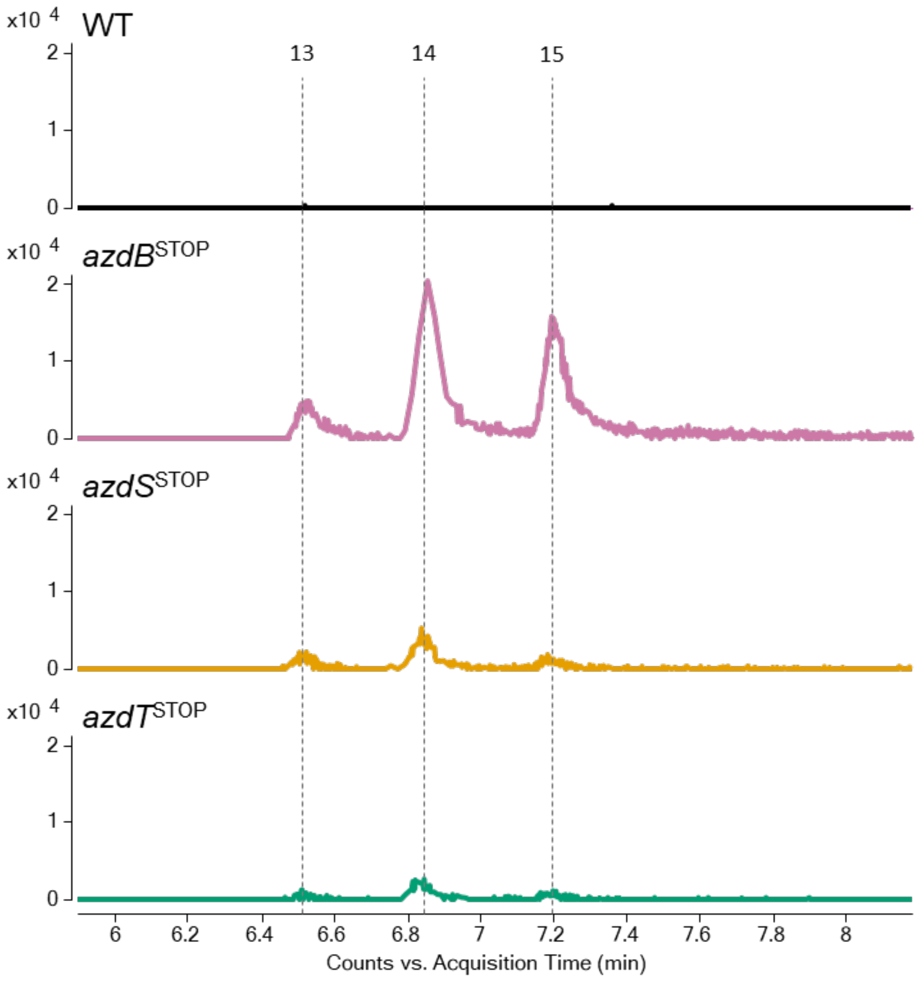

Figure S.20: Extracted ion chromatogram of compound **13-15**, where the first and the last eluting peak correspond to the smallest and the largest EIC respectively.

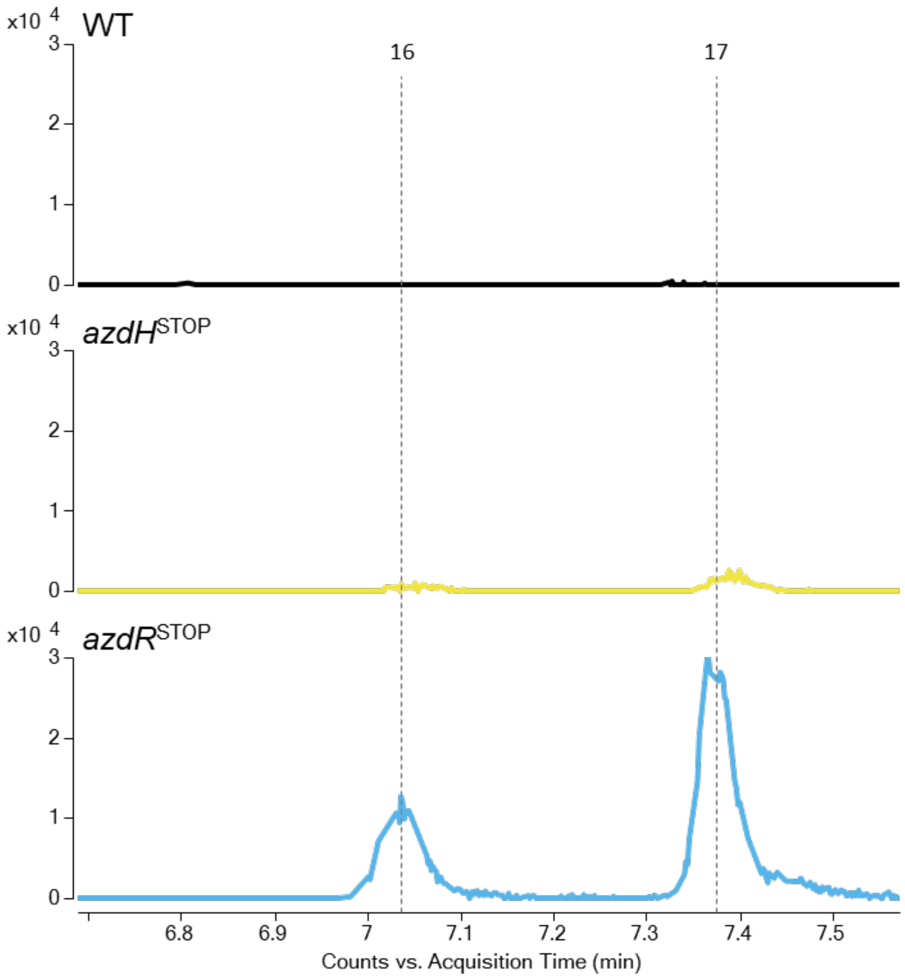

Figure S.21: Extracted ion chromatogram of compound **16-17**, where the first and the second eluting peak correspond to the smallest and the largest EIC respectively.

### Insights into azodtrecin-like BGCs

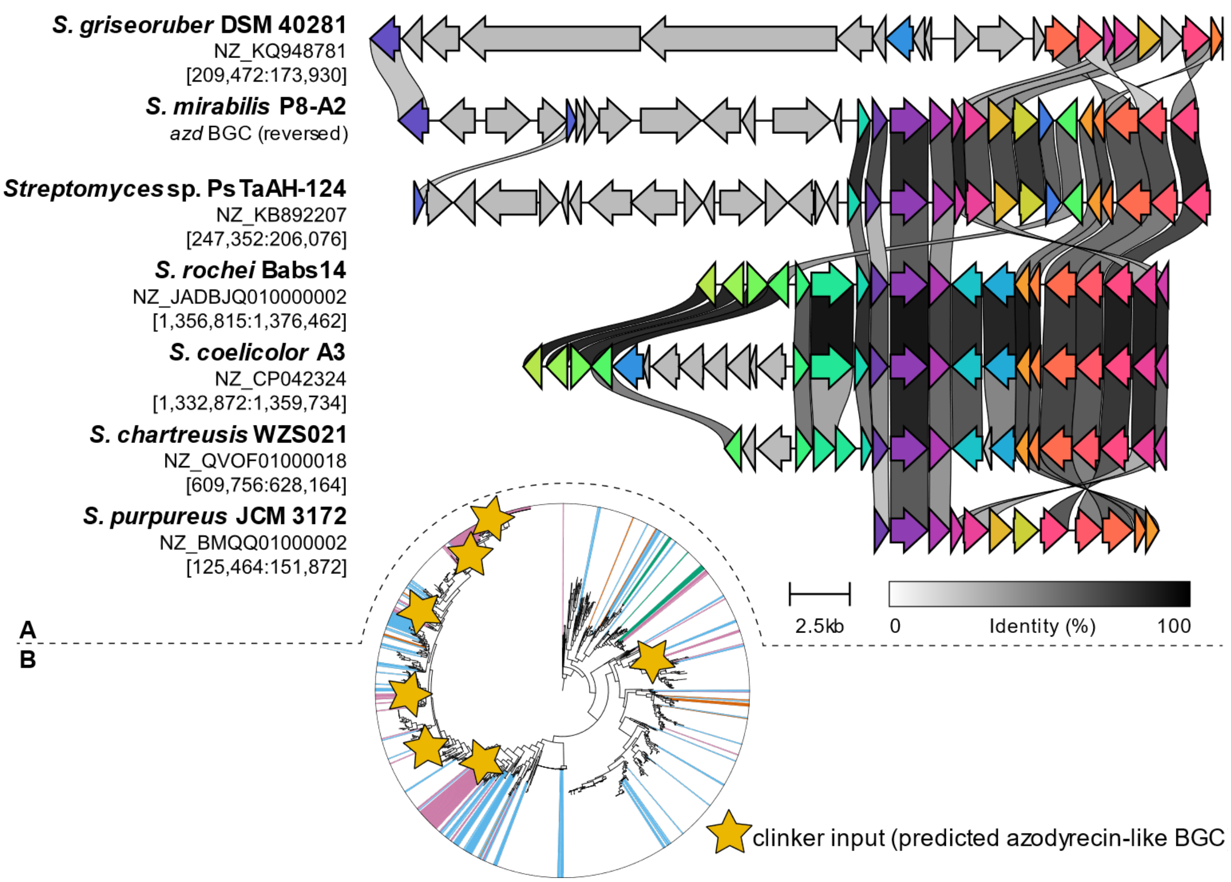

Figure S.22: Comparison of phylogenetically distant biosynthetic gene clusters detected by GeneClusterPhyloMapper. (**A**) A random set, including S. coelicolor A3 and azodyrecin producer S. mirabilis P8-A2, of predicted BGCs were compared using clinker[46]. (**B**) overview of phylogenetic distance between input BGCs visualized on Streptomyces whole genome phylogenetic tree generated using autoMLST, enlarged tree available in Figure 7.
